# Anatomics MLT, an AI tool for large scale quantification of ultrastructural traits

**DOI:** 10.1101/2024.05.06.592763

**Authors:** Aaron Brookhouse, Yuting Ji, Jan Knoblauch, Petr Gaburak, Brittney M. Wager, Zudi Lin, Hans-Henning Kunz, Assefaw H. Gebremedhin, Hanspeter Pfister, Winfried S. Peters, Donglai Wei, Michael Knoblauch

## Abstract

The ever increasing breadth of biological knowledge has led to recent efforts to combine information from various fields into cell– or tissue atlases. Anatomical features are the structural basis for such efforts, but unfortunately large scale analysis of subcellular anatomical traits is currently a missing feature. Similarly, small phenotypic alterations of organelle– or cell-specific anatomical traits, such as an increase of the total volume or the number of mitochondria in response to certain stimuli, are currently hard to quantify. To provide tools to extract quantitative information from available 3D microscopic datasets generated with methods such as serial block face scanning electron microscopy we a) developed much improved fixation and embedding protocols for plants to drastically reduce processing artifacts and b) generated an easy-to-use AI tool for quantitative analysis and visualization of large-scale data sets. We make this tool available as open source.

## Introduction

The quantification of anatomical data is of increasing importance in biology. Currently, many phenotypic traits remain undiscovered due to a lack of appropriate methodologies to detect small cellular and subcellular alterations. The identification of, e.g., a slight reduction of cell wall thickness in a specific cell type, or a small increase of the number or total volume of mitochondria in a tissue layer currently are not possible without immense effort. Other efforts including the generation of cell or tissue level maps require detailed quantitative anatomical data. Recent developments in 3D reconstruction at the cellular level after serial imaging by confocal microscopy has facilitated progress in the identification of changes in cell shape and pattern in specific tissues (Wolny *et al*., 2020). The resolution of light microscopes, however, is limited, and detailed 3D information on subcellular structures has to be acquired by electron microscopy (Kievitz *et al*., 2022).

The classical approach to 3D reconstruction of cellular and subcellular structures by transmission electron microscopy (TEM) is serial sectioning followed by the reconstruction of 3D relationships from the stacks of 2D micrographs (Bang & Bang, 1957; Harris *et al*., 2006; Hoffpauir *et al*., 2007; for a modification based on scanning electron microscopy [SEM], see Horstmann *et al*., 2012). However, these methods are comparatively labor-intensive and the handling of large numbers of sections can be difficult. An alternative for small objects such as organelles or protein complexes is electron tomography, in which the object is imaged while being tilted in numerous small steps (Ishikawa, 2016; Otegui, 2020). To acquire large scale tissue-level information from hundreds or thousands of cells while still allowing sufficient resolution for the identification of structures such as golgi, plasmodesmata, or nuclear pores, serial block face scanning electron microscopy (SBFSEM) has been developed. With this technique, the object is sectioned by a microtome located within an SEM. After an early proof–of-principle by Stephen Leighton (1981), essential improvements were accomplished by Denk & Horstmann (2004), which turned SBFSEM into the method of choice in diverse fields (for reviews, see Hughes *et al*., 2014; Zankel *et al*., 2014; Kremer *et al*., 2015; Cocks *et al*., 2018; Smith & Starborg, 2019, Knoblauch *et al*. 2024). The plane surface of a resin-embedded, appropriately stained sample, or block, is imaged utilizing backscattered electrons, and the block is advanced by a set distance. Then, a thin section is removed by an ultramicrotome, or by milling with a focused ion beam (Peddie & Collinson, 2014). Thus a new block face is revealed, and the process repeats. Since not the section but the block face is imaged, compression distortions, which are common in ultrathin sections, usually do not pose problems. Issues arising due to insufficient conductivity of the block often can be avoided by applying heavy metal stains, or can be mitigated by surrounding the specimen with metal-impregnated resin (Wanner *et al*., 2016) and by focal gas injection (Deerinck *et al*., 2018).

Electron microscopy provides much greater resolution than light microscopy, but does not allow for the observation of living material, as samples need to be fixed, dehydrated, stained, and in many cases subjected to contrast-enhancing treatments. These treatments can drastically affect the appearance of subcellular structures such as plasmodesmata (Peters *et al*., 2021). Sample vitrification by rapid cryofixation under high pressure can avoid some of the artifacts potentially caused by chemical fixation, but the maximum sample size up to which this method provides satisfactory results is limited (Dubochet, 2007; McDonald, 2009). In our hands, cryofixation does not work well with samples of plant tissues consisting of multiple layers of mature cells.

A protocol based on the standard fixation procdure originally introduced in 2010 for the SBFSEM study of mammalian tissues (Deerinck *et al*., 2022) soon became adopted by plant scientists. This protocol, however, tends to produce artifacts in plant tissues such as collapsed, undulating cell walls and star-shaped vacuoles. In more or less substantially modified variants, the Deerinck method was applied, for example, to reconstruct developing sieve plates in the phloem (Dettmer *et al*., 2014) and unusually shaped nuclei in carnivorous plants (Płachno *et al*., 2017), to determine cell and chloroplast volumes at increased precision (Harwood *et al*., 2019), and to produce 3D visualizations of developing chloroplasts (Pipitone *et al*., 2021). All segmentations and quantitative analyses were performed on one or a few cells. Lippens *et al*. (2019) adopted the protocol to obtain 3D visualizations of *Arabidopsis* root tips, and segmented plasmodesmata in a selected region after intense thresholding. So far, however, large-scale quantitative anatomical analyses in plants are lacking.

As emphasized already by Leighton (1981), the value of SBFSEM will be greatest when full automation is achieved, and large numbers of digital images can be processed by appropriate software. Some of the required tools are now available, and commercial as well as open-source software for different tasks has been developed (Hughes *et al*., 2014; Kittelmann *et al*., 2016; Cocks *et al*., 2018). Compared to classical serial sectioning, registration (the correct alignment of the images of a z-stack) is less of a problem in SBFSEM since the block does not move laterally during the procedure, and image alignment is a standard procedure in SBFSEM instruments. On the other hand, image segmentation – the partitioning of an image into segments representing, for example, different organelles in an imaged cell – still is far from being fully automated. Due to the necessarily varying staining conditions of tissue samples optimized for the visualization of different structures, traditional image analysis methods may generate results that require a significant amount of manual labor for error correction. Recently, deep learning methodologies have shown great success in segmenting electron microscopy images, which requires mask annotation and model training on a relatively small number of images (Lee *et al*., 2017, Januszewski *et al*., 2018, Sheridan *et al*., 2023).

We aimed to close remaining methodological gaps by providing a comprehensive toolbox for extracting quantitative anatomical data from hundreds or thousands of cells by SBFSEM. First, we modified fixation and staining protocols based on the current standard methodology by Deerinck *et al*. (2022) for improved contrasting of cellular structures specifically in plant tissues, a critical step to render image stacks suitable for analysis by deep learning algorithms. In addition, the new protocols strongly reduced artifacts in plant tissues. Second, we developed Anatomics MLT, a software package based on the open-source deep learning segmentation software PyTorch Connectomics (PyTC; Lin *et al*., 2021), for large-scale analysis and optimized reliability of the automated identification of specific types of objects in image stacks.

## Methodology

### Preparation protocols for serial block-face scanning electron microscopy (SBFSEM)

An osmotically induced, intracellular hydrostatic or turgor pressure that approaches or even exceeds 1 MPa is a biophysical feature that distinguishes the walled cells of plants from animal cells, which lack cell walls and therefore cannot sustain significant intracellular pressure (Peters *et al*., 2000). Turgor pressure in plant cells stretches the cell walls. This turgor-induced elastic cell wall strain can be substantial. In growing roots, an extreme example, linear elastic strain may reach 0.3 in the tip meristem (own unpublished results). Notably, preparation for electron microscopy necessarily includes dehydration of the sample, which induces a loss of turgor pressure even before the cells are sectioned. Therefore, EM preparation protocols optimized for unpressurized animal cells rarely yield satisfying results when applied to turgescent plant tissues.

Initially, we used Arabidopsis root tips to test over 130 different protocols to reduce artifacts and optimize staining for specific organelles. Figure 1 shows some of the artifacts induced in an *Arabidopsis* root prepared by the standard protocol for animal tissues (Figure 1 A,B; Deerinck *et al*., 2022). The walls appear folded and undulated (cytorrhysis) and vacuoles are collapsed. Both shrunken vacuoles and cytorrhysis are symptoms of significant cellular volume loss. Collapsed vesicles and the wavy appearance of the nuclear envelopes support the interpretation that the cells had experienced significant volume losses by the time their internal structures became fixed. These effects appear particularly unacceptable when they occur with methods that aim specifically at 3D, *i.e.*, volumetric reconstructions of tissues and cells. Following a series of experiments, we were able to identify modifications of the protocol that avoid or at least significantly reduce these artefacts, while providing strong contrast for the analysis of different organelles.

**FIGURE 1.**
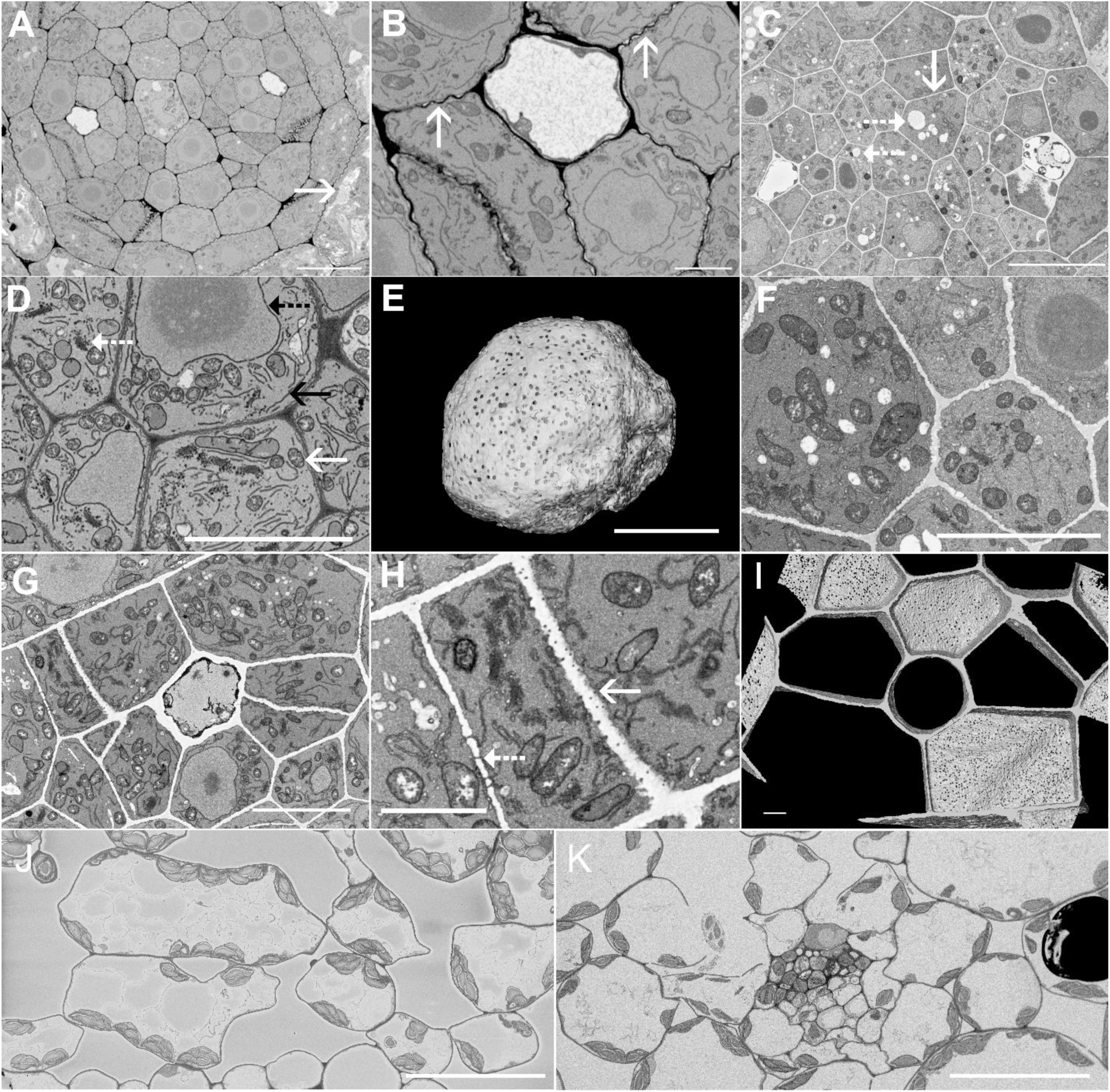
Optimization of protocols for plant tissues. The standard SBFSEM preparation protocol developed for animal tissues (Deerinck et al., 2022) leads to artifacts when applied to plant material, demonstrated here in cross-sections of the central cylinder of an Arabidopsis root tip (A,B). Vacuoles are collapsed (arrow in A) and the cell walls, which appear dark, are wavy, typical of cytorrhysis due to volume loss of the protoplast (arrows in B). Modifications of the protocol, especially the adjustment of the osmolarity of the fixation medium, the fixation schedule, dehydration, and resin embedding procedures (see main text) lead to straight cell walls, and to an inflated appearance of vacuoles and vesicles (C). Application of standard stains over altered durations compared to the standard animal protocol results in well-stained membranes and cell walls (D). Nuclear membranes (dashed black arrow), ER membranes (solid black arrow), mitochondria (solid white arrow), and Golgi (dashed white arrow) are well visible (D) and 3D reconstructions of individual organelles such as the nucleus with nuclear pores are possible (E). Alteration of the standard plant protocol by exchanging potassium ferrocyanide for potassium ferricyanide reduces the staining of cell walls and membranes while increasing that of the mitochondrial lumen, which facilitates automated recognition of this organelle (F). Omission of uranyl acetate and Walton’s lead in the staining procedure results in unstained cell walls (G,H). Due to the staining of membranes, plasmodesmata crossing the cell walls appear very well contrasted (solid arrow in H shows cross-sectioned plasmodesmata, dashed arrow highlights longitudinally sectioned plasmodesmata). A 3D reconstruction of the cell wall visualizes plasmodesma density and distribution (I). Osmotic effects of the fixation media on cell preservation can be significant, especially in thin-walled cells that are facing air spaces, like in the mesophyll (J,K). Cells in J were fixed at higher osmotic potential (100 mM cacodylate buffer) than in K (25 mM cacodylate buffer). In J, cell structure is well preserved. In K, cell walls and some organelles are ruptured, and some organelles are found in extracellular spaces. Scale bars: A, 20 μm; B, D, I, F, 5 μm; C, 10 μm C; E, G, 2 μm; H, 1 μm; J, K, 30 μm.

*Fixation* — Fixatives of high osmotic potential (isotonic or hypertonic) prevent inflation and potential bursting of animal cells. In contrast, a hypotonic fixative is beneficial to maintain pressure and reduce shrinkage in plant cells. Compared to the standard protocol, we omitted paraformaldehyde while slightly increasing the concentration of glutaraldehyde for fixation. Most importantly, we significantly reduced the buffer concentration from 150 mM to 25 mM. Because comparably large tissue fragments need to be embedded and evenly fixed, the incubation period was doubled to allow diffusion of the fixative to the innermost parts of the sample. Finally, once the fixative reached all parts of the sample, we exposed the tissue to microwave radiation. Microwaves drastically accelerate the diffusion of polar molecules and can aid fixation within minutes or seconds. However, it is crucial that microwaves are applied only after the fixative has reached all parts of the sample, to prevent microwave-induced structural artifacts. The addition of a microwave treatment following the initial incubation significantly improved the fixation, especially at a distance from thespecimen surface. Washing with ddH_2_O rather than buffer seemed to improve results. Figure 1 C shows an overview of a root tip following the above described treatment. The cell walls are straight and cells appear turgescent. Vacuoles and other organelles remain round and do not collapse.

*Staining* — Staining times were extended in all three post-fixation steps to achieve even staining as plant cell walls strongly reduce diffusion compared to animal tissues. Exposure to osmium tetroxide (OsO_4_) in the presence of potassium ferrocyanide (K_4_Fe(CN)_6_) followed by thiocarbohydrazide (TCH) and OsO_4_ again provided adequately contrasted plant membranes for SBFSEM imaging. Decreasing the OsO_4_ concentration in the first step provided improved results. As ferrocyanide strongly stains membranes and other structures such as the cell wall, we omitted additional staining with uranyl acetate and Walton’s lead. The resulting images (Figure 1 D) allow clear identification of ER membranes, mitochondria, nuclear membranes, golgi, and other intracellular structures.

*Dehydration* — Best results were obtained by applying acetone rather than ethanol series. Stepwise dehydration in 30%, 40%, 50%, 60%, 70%, 80% 90% acetone followed by three exchanges in 100% at room temperature worked well.

*Resin embedding* — We produced better results with hard Spurr’s resin than with the commonly used Durcupan ACM resin. As Spurr’s resin polymerization is temperature-dependent, slow polymerization occurs at room temperature which increases the viscosity over time. Larger samples may show insufficient infiltration in the central parts. To accommodate longer infiltration times, omission of the accelerator, DMAE, from the standard Spurr’s recipe for the graded infiltration series (Spurr, 1969) may be applied. Inclusion of DMAE only in the final three 100% infiltration steps (including the final microwave treatment) facilitates proper polymerization even in central parts of the sample.

*Imaging* — The high density of the cytoplasm and the small size of vacuoles makes imaging of root tips in the high-vacuum mode prepared as above unproblematic. This is due to high concentrations of heavy metals (stains) that mitigate charging. In tissues that contain large, practically unstained vacuoles, charging of the sample can occur, which may require imaging in the low vacuum mode to prevent image artifacts. The same is true for leaf tissues with large intercellular spaces. Imaging in low vacuum mode reduces the achievable resolution which may complicate the identification of very small objects such as plasmodesmata, but organelles such as mitochondria remain well identifiable.

Application of this Standard Plant Protocol (Protocol 1; Table 1) provides satisfying results with turgescent tissues of *Arabidopsis* (Figure 1D). Cell walls appear straight as if still under tension. Vacuoles, vesicles, organelles are round and mostly without indentations, and the undulation of the cell wall and nuclear membrane is mitigated. The cytoplasm seems dense and cytorrhysis is not observed, even in the central cylinder. Delicate structures such as the Golgi are resolved. Well-stained membranes allow for 3D reconstruction of structures such as the nucleus highlighting the number of nuclear pores (Figure 1 E). In the following, we describe modifications of this Standard Plant Protocol for specific purposes.

**Table 1.** Fixation and Embedding Protocols.

| **Table 1. Fixation and Embedding Protocols** |  |  |
| --- | --- | --- |
| **Protocol 1** (Standard Plant Protocol) | Time | Alteration for alternative protocols |
| Fixative:<br>4% GA, 0.025M Cacodylate, pH 6.8, 2 mM CaCl<sub>2</sub>, room temperature | 6 h |  |
| Leave in fixative and fix in microwave at 300W, 35°C cutoff | 2 min |  |
| rest at room temperature | 5 min |  |
| Rinse with 0.025 M Cacodylate | 3 × 10 min |  |
| Apply 1.5% K<sub>4</sub>Fe(CN)<sub>6</sub>, 0.025M Cacodylate, pH 6.8, 2 mM CaCl<sub>2</sub>, 2% OsO<sub>4</sub>, 4°C, *Mix before adding* | over night | Exchange K<sub>4</sub>Fe(CN)<sub>6</sub> for K<sub>3</sub>Fe(CN)<sub>6</sub> in **Protocols 2 and 3** |
| Rinse with dd water | 3 × 10 min |  |
| 0.5% TCH *Make fresh; takes ~ 30 to 45 min to dissolve in 60 °C oven. Cool to RT and filter before use.* | 45 min |  |
| Rinse with ddH<sub>2</sub>O | 3 × 10 min |  |
| 2% OsO<sub>4</sub> | 3 h |  |
| Rinse with ddH<sub>2</sub>O | 3 × 10 min |  |
|  |  | **Protocol 3:** 2% uranyl acetate over night at 4° C, 3 × ddH<sub>2</sub>O rinse 10 min each, Walton's lead for 60 min at 60°C 3 × ddH<sub>2</sub>O rinse 10 min each. |
| Dehydrate in 30%, 40%, 50%, 60%, 70%, 80%, 90%, 100%, 100%, 100% acetone | 10 min each |  |
| Acetone: hard Spurr's resin without DMAE 3:1, 2:1, 1:1, 1:2, 1:3, 100% hard Spurr's without DMAE | 8-24 h each |  |
| 2 × 100% hard Spurr's resin with DMAE | 24 h each |  |
| Microwave in 100% hard Spurr's (with DMAE) at 100 W, 40°C max | 60 min |  |
| Cure at 70°C | 48 h |  |

### Optimization of protocols for specific cell types/ structures

Increased contrast of the structure of interest relative to the surrounding material is the main factor in improving the automated recognition of cellular components (see below). Therefore we modified staining protocols to increase contrast for specific structures.

*Post-fixation option: ferrocyanide* vs. *ferricyanide **(Protocol 2)*** — In plant and animal cells, staining with OsO_4_ combined with potassium ferrocyanide, K_4_Fe(CN)_6,_ provides good overall contrast of membrane systems and is therefore commonly applied for SBFSEM (Lippens *et al*., 2019). However, it tends to stain cell walls whereas potassium ferricyanide, K_3_Fe(CN)_6_, does not (Figure 1 F). Unstained cell walls are especially important when observing plasmodesmata. The strongly stained desmotubule and plasmamembrane provides excellent contrast to reconstruct cell walls including plasmodesmata (Figure 1G, Protocol 2).

*Uranyl acetate and Walton’s lead **(Protocol 3)*** — Uranyl acetate staining and subsequent incubation with Walton’s lead aspartate improves the conductivity of the block and further enhances overall contrast. In combination with ferricyanide, we found the staining especially suitable to observe mitochondria as their lumen becomes well contrasted (Figure 1 I). This protocol is also suitable for large-scale reconstruction of vacuoles, as uranyl acetate and Walton’s lead produce an increased staining of the cytoplasm and the cell walls, while vacuoles remain unstained (Figure 1 I, Protocol 3).

*Arabidopsis leaf tissue **(Protocol 4)*** — Leaf tissues – especially the mesophyll – are very delicate as cell walls are thin and many cells do not share walls with neighboring cells because they face intracellular air spaces. Since osmotic effects due to shifting rates of photosynthetic sugar production are common, a critical factor in preserving intracellular structures in leaf tissues is the osmolarity of the fixative, which can be most easily adjusted by changing the buffer concentration. Low buffer concentrations that are beneficial for maintaining cell structure in root tips may lead to artifacts in leaves. Figure 1 J shows *Arabidopsis* leaf mesophyll cells after processing with a fixative containing 100 mM cacodylate buffer; the overall structure is well preserved. When treating the same tissue with equal concentrations of fixatives but only 25 mM buffer, cell walls are ruptured and organelles are released into extracellular spaces (Figure 1 K). As osmolarities especially in leaves may change quickly depending on light and water availability, buffer concentrations need to be adjusted accordingly. The plants used in this study were harvested in the afternoon, resulting in large starch grains in the chloroplasts.

## Automated image analysis

### A) Label generation

Over the years, several software packages have been developed that have, despite their potential usefulness, resulted in limited usage by biologists because of the requirements of programming skills that many researchers are lacking. In addition, many commercial software packages are cost-prohibitive for a majority of labs. Our intention was to create a free and easy-to-use software package that can be operated by simple mouse clicks instead of program language input into command prompts. In the following, we will highlight key features of Anatomics MLT that are critical to gain quantitative results with high precision as well as 3D image reconstruction. The full documentation explaining all possible input features is provided as an Appendix below and online under https://pytorchconnectomics.github.io/Anatomics-MLT.

Prerequisite to the training of the software for the identification of specific structures such as organelles is the generation of labels, copies of selected images from the stack on which the structure(s) of interest are highlighted and everything else is masked as background (Figure 2). The number of labels required depends on how distinct the structure of interest is. The speed and efficiency of the training procedure therefore depends on the appropriateness and quality of the staining that had been applied to the samples before image acquisition. Labels can be generated with any image processing software including ImageJ, Blender, Photoshop, and others. The label background must be black (grayscale value 0), otherwise the software will recognize it as a feature of interest. If the structure of interest is sufficiently distinct, standard image processing tools such as thresholding or selection by “magic wand” and subsequent filling can be used. In the worst case, features will have to be labeled manually to generate a mask that perfectly matches the structures of interest (Figure 2 B,C). Assuming that an imaging stack of 1000 images has been generated, a good starting point is to generate 20 labels from consecutive images, which will have to be saved as an image stack in a single TIF file (see imaging tools in our software, or open ImageJ > file, import, image sequence > import all 20 images > file, safe as > choose tiff and save). Another image stack consistring of the 20 original images used to generate the labels also needs to be generated. The two stacks will serve as input training files for the software.

**Figure 2:**
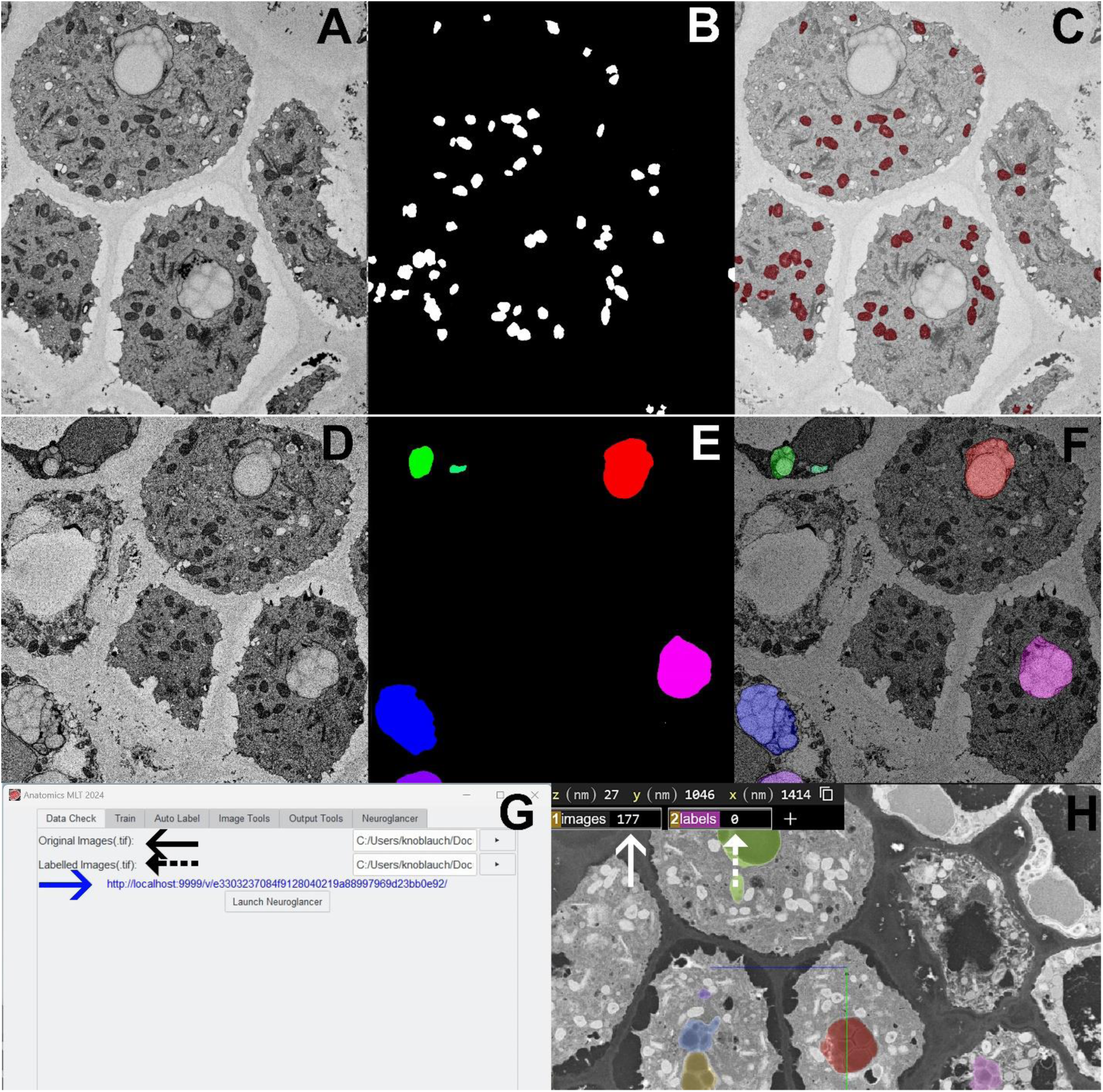
Generation of Labels for Anatomics MLT. A) Single slice of a 3D stack of the columella in the root tip of Arabidopsis using our modified Protocol 3 B) generation of a “Semantic Label” (white: structures of interest; black: background) that precisely identifies all mitochondria in the section, informing the algorithm of the location of the structures of interest. C) Overlay of (A) and (B). D) Different image in the same 3D stack as in (A). E) An “Instance Label” in the columella highlights each individual statolith in a different color. F) Overlay of (D) and (E).G) The “Data Check” pane in Anatomics MLT allows to check for labeling accuracy. H) The Neuroglancer software running in a web browser enables dimple double-checking of label values (see text for details).

Anatomics MLT works with two different label types. In “Semantic Labeling”, all individual structures of a certain type (e.g. all mitochondria in the image) are labeled in the same color (typically white, Figure 2 B). In “Instance Labeling”, individual occurrences of the same structure of interest are labeled in different colors (Figure 2 E). Instance labeling has the advantage that individual structures that are close to each other, or even touch each other, labeling each individual organelle as a separate unit assists the algorithm in recognizing that the structures of interest should not be treated as a single large complex. However, in instance processing only one structure of interest can be processed at the same time. Semantic labeling on the other hand allows for the labeling of multiple structures such as mitochondria and the nucleus at the same time. In this case the background of the labels would be black and all mitochondria would be labeled in one color (e.g. white) and all nuclei in a different color (e.g. red).

The software has 6 main panes. The first is the “Data Check” pane (Figure 2 G,H), which we included for users who are unfamiliar with more advanced imaging software. To verify that stacks and labels are accurate, the “Data Check” pane is opened and the image stack (black arrow) label stack (dashed black arrow) are loaded. Clicking “Launch Neuroglancer” opens a link (blue arrow) that will open the Neuroglancer software in the default browser of the computer used. Neuroglancer will show the images and labels as overlays, enabling visual inspection of their accuracy (Figure 2 H). In the upper left, numerical grayscale values of the pixels under the cursor on the original image (white arrow) and on the label (dashed white arrow) will be seen. The latter must be 0 when moving the cursor over a background area, else the background will be identified as an object of interest by Anatomics MLT. With the cursor over a labeled area (any colored area), the value must be 1 or higher. For semantic labels, all label values will be the same. In the case of instance labels (various colors as in H), values will differ and Anatomics MLT will distinguish individual instances.

### B) Training the Software

Figure 3 shows the individual panes of Anatomics MLT. First, the generated image stack (Figure 3, solid black arrow) and label stack (dashed black arrow) are loaded into Anatomics MLT. Next, the x,y,z, voxel dimensions (i.e., the voxel resolution in 3D, for example: 10 nm x, 10 nm y, 40 nm z) have to be defined (dashed blue arrow). If the x, y, and z dimensions are equal, values can be set to 1 as the program processes relative dimensions only. The quality of the training and hence also that of the output data is defined by multiple parameters. The number of iterations (Figure 3A, solid blue arrow) defines how many times the training is repeated. Usually, higher numbers improve quality, but may also lead to significantly longer computing times. Larger numbers of labels may be needed if the contrast between structures of interest and background is low. The window size (red arrow in Figure 3A) indicates the voxel dimensions of the 3D window (x, y, z) that the software utilizes to identify the defining features of the structure of interest. It is important that the window’s x and y size exceed the size of individual structures to be identified. For instance, if the size of a nucleus is in the range of 200 × 200 × 200 voxel, a window size of 150 × 150 × 150 would produce poor results. In this case a window size of 250 × 250 × 250 voxel at minimum needs to be used. Thus, small structures such as mitochondria require smaller window sizes which reduces computing time. Increasing window size for small structures would not be of benefit. Once the algorithm is trained, it may be applied to various image stacks as long as contrast and shape of the structure of interest are similar.

**Figure 3:**
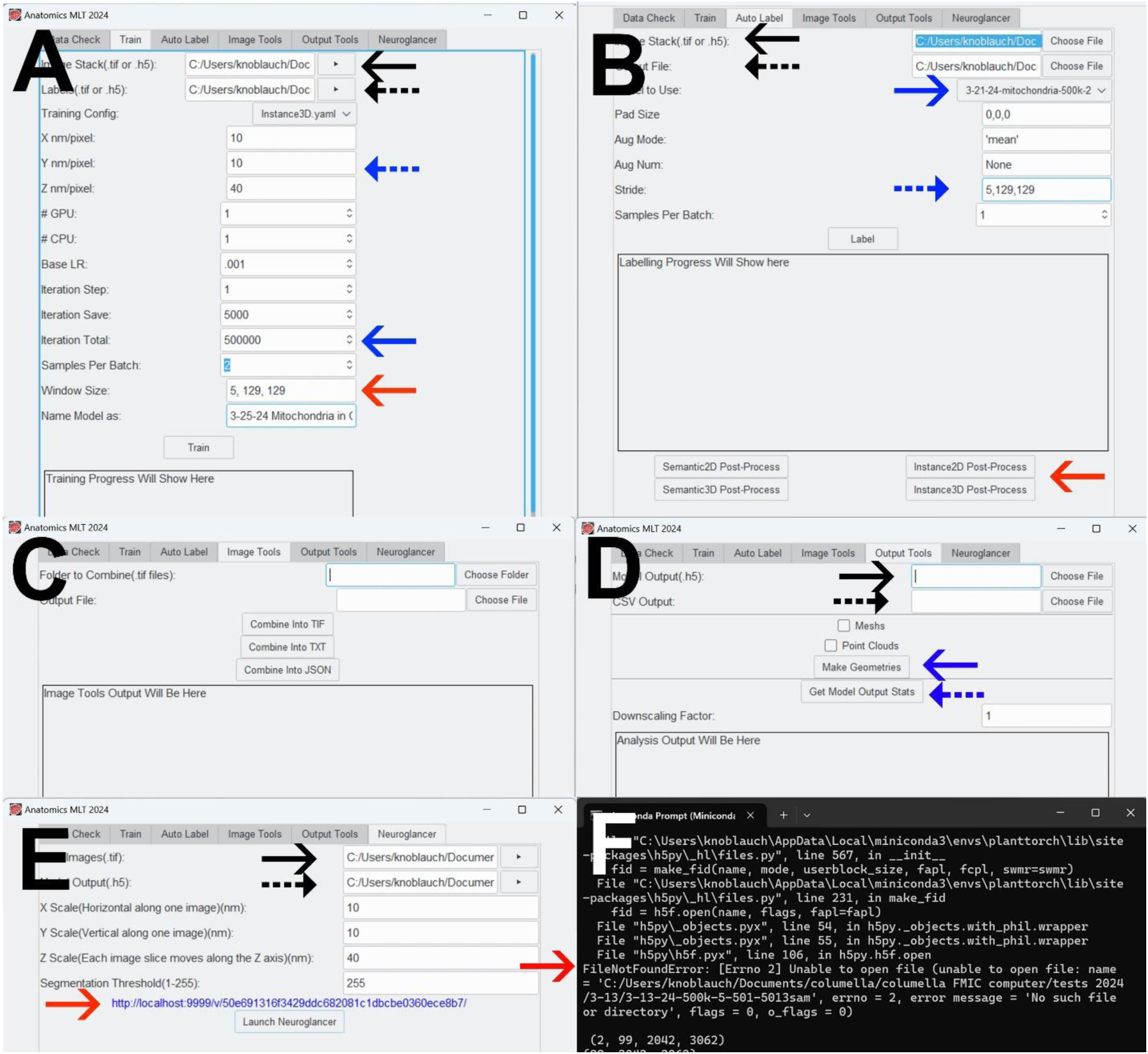
Anatomics MLT Software. A) Key features of the “training pane” of Anatomics MLT are the input window of the training stack (black solid arrow), the labels stack (black dashed arrow), the x,y,z dimensions (blue dashed arrow), the number of iterations which significantly impact the training quality (solid blue arrow) and the training window size (red arrow). Other adjustable parameters have less impact on overall quality and are explained in the appendix. B) The “auto-label” pane uses the generated training file to identify the structure of interest throughout the whole stack. Key input data are the full image stack (black solid arrow), a chosen file name for the final output file (black dashed arrow), the choice of the appropriate training file from the drop down menu (solid blue arrow), and defining the stride (blue dashed arrow; see main text). In many cases a post-processing step will be required (solid red arrow) which, if necessary, will be indicated in the text box. C) The “image tools” pane allows the combination of individual images into stacks of different format. D) the “output-tools” pane allows for the extraction of different data files. Choosing the output model file (.h5 file) generated during auto-label (solid black arrow) and clicking “make geometries” (solid blue arrow) will generate a new mesh file and a new point cloud file. Those files can be imported into other software packages like Blender or Avizo for 3D visualization. Choosing a file name (dashed black arrow) and clicking “get model output stats” (dashed blue arrow) will generate an excel CSV file that includes all incidents (e.g. all individual target organelles), their individual volume, the total volume of the stack, etc. E) The “Neuroglancer” pane allows the 3D visualization of the generated data in the Web-based Neuroglancer viewer generated by the Google Connectomics Team. Input of the raw image stack (solid black arrow), the output model file generated during auto-label (dashed black arrow), and clicking “launch neuroglancer” will create a link (red arrow). Clicking the link will launch the default web browser and display the generated 3D reconstruction. F) The anaconda prompt may provide important information, e.g. when errors occur. For example, storing the original 3D stack in a different folder than the output file creates an error (solid red arrow) when clicking the post processing button. In this case an output file (dashed black arrow in B) needs to be redefined.

### C) Auto-Label

Once the algorithm has been trained, the training file is used to auto-label the entire image stack in the auto-label pane (Figure 3 B). For this purpose, either a subset or the full stack of the original reconstruction is chosen, but the subset of images that was used for training should not beincluded. The image stack is loaded into the first box (solid black arrow), a name for the file to be generated is defined (dashed black arrow) and the previously generated training file is selected by clicking the drop down menu (solid blue arrow). The final parameter in the auto label pane of major importance is the stride (dashed blue arrow). The algorithm does not process the entire image stack at once but works on parts of it at a time. After it has finished with such a section, it moves on to process another section of the same size. The distance it moves after processing one section is the stride. If the stride is the exact same size as the windows size used during training, each section will be processed once. The stride should not be bigger than the window size, as otherwise certain sections will not be processed. If it is smaller, some parts will be processed repeatedly. Clicking “label” will start the process and will generate an .h5 file.

In many cases a post-processing step is required which will be indicated in the text box (dashed red arrow). If, e.g., “instance 3D post-processing” is indicated in the text box, click this button (red arrow) and a new final .h5 file will be generated. Any errors that may have occurred will be listed in the Anaconda Prompt (see Figure 3 F).

### D) Image Tools

The fourth pane provides tools to generate 3D stacks from individual images (Figure 3 C).

### E) Output Tools

In the Output Tools pane (Figure 3 C), choosing the .h5 file (solid black arrow) that was generated by the auto-label process and clicking “make geometries” will create either a mesh file or a point cloud file or both, depending on which boxes are checked. The generated files can then be loaded into programs such as Blender or Avizo for 3D visualization.

The output tools pane also has the option to extract quantitative data. Again, the .h5 file is loaded (solid black arrow), an output file name is chosen (dashed black arrow) and “get model output stats” is selected. This will create a .csv file that contains all individual instance IDs (all individual organelles of interest) and indicates the volume of each of the structures. In addition, the total volume of the stack is indicated allowing for statistical analysis.

### E) Neuroglancer

Although Anatomics MLT can generate point clouds and meshes for import into various image processing tools, we created a pane that allows direct access to Neuroglancer (Figure 3 E), a high-performance, flexible WebGL-based viewer and visualization framework for volumetric data developed by the Google Connectomics Team. Although initially created for neuroscience applications, Neuroglancer is highly useful in various contexts. Here, we provide a brief tutorial on how to obtain 2D and 3D images.

For Neuroglancer visualization, the image stack (raw images, solid black arrow in Figure 3E) and .h5 file generated by the auto label procedure (model output, dashed black arrow in Figure 3E) need to be defined. After input of the dimensions etc., the box “launch Neuroglancer” is clicked and a link (red arrow in Figure 3E) will appear. This may take a few minutes depending on file sizes. Selecting this link will open the default web browser on the computer (Figure 4) and will display the reconstruction in 4 panels, x, y, z, and xyz (Figure 4 A). Figures 4 B-D show images of columella cells from the same stack, without labels (B), statoliths labeled (C), and mitochondria labeled (D). Users can scroll through the stack in the z direction. To obtain 3D images, the layer side panel button (blue dashed arrow in E1) and then the Edit JSON button (solid blue arrow in E1) have to be clicked. A window will open (E2), in which “Selected layer” is opened by clicking the arrow next to it (black dashed arrow). The word “image” needs to be replaced by “gt” (ground truth; black solid arrow), and “apply changes” and “close” are clicked to apply the changes. “Seg” will now appear in the layer panel (white dashed arrow in E3). This selection will open a new pane. In a previous step (under output tools in the Anatomics MLT software), a CSV file was generated containing all identified instances. This file contains three columns. The second column is selected and pasted into the “select” window (white solid arrow in E3). All individual instances (e.g. individual identified organelles) will now be listed. Highlighting an instance by clicking the box next to it will display the 3D shape of the specific structure. Depending on file sizes and the quality of the internet connection used, these data visualizations may require some time.

**Figure 4:**
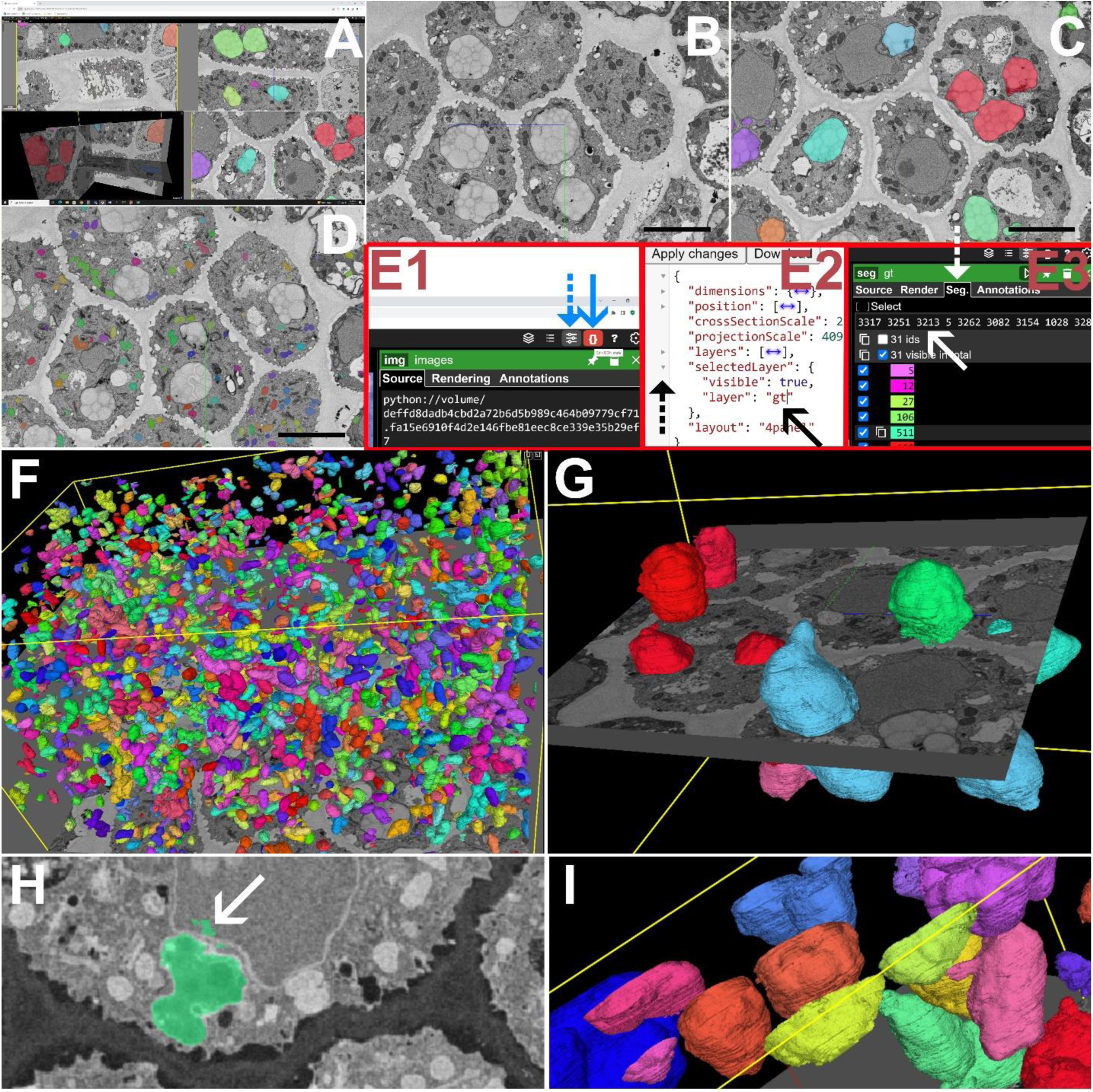
Using Neuroglancer for visualization and statistics. A) The standard neuroglancer surface provides x, y, z, and xyz views of an image stack. View of the x plane (B) with highlighted statoliths (C) or mitochondria (D). The generation of a 3D view of the organelles of interest requires a few steps (E1, E2, E3) that are described in the main text. F) 3D visualization of all ∼2800 mitochondria in a columella reconstruction. G) Selected statoliths in the columella reconstruction. H) Identification errors (arrow) that may occur depending on the quality and number of labels, training iterations, etc, can be manually eliminated by a double click on the structure, which makes it disappear. The same can be done with partial structures that are located at the margins of the 3D reconstruction (I). Scale Bars: B,C,D, 5 µm.

Figure 4 F shows all mitochondria in the volume of the columella. 2862 individual mitochondria have been identified with a total volume fraction of 3.75%. Figure 4 G highlights a few selected statoliths with an average volume of 43.9 µm^3^ (sd ± 19.3 µm^3^). For more data that have been processed using Anatomics MLT please see Knoblauch et al. (2024).

### F) Elimination of errors

Misidentifications by the software may occur (Figure 4 H, arrow), especially where structures of interest are cropped close to the borders of the stack. Misidentifications may be eliminated manually by identifying the instance number and deleting it. This seems reasonable if only a few structures of interest are present. If misidentifications have implausibly small volumes, a reasonable volume cutoff may eliminate all smaller structures. If too many misidentifications are present, new training with more labels and adjustment of parameters such as the number of iterations may be required to improve quality. What strategy is least laborious depends largely on the number of structures of interest contained in the stack volume. Manual elimination appears too labor-intensive in the case of mitochondria in the columella (Figure 4 F), while it seems reasonable in case of statoliths in the columella. In addition, CSV files contain the volumes of all partial structures that extend beyond the borders of the stack (e.g. Figure 4 I). These create an error that increases with decreasing stack volume. Therefore manual elimination and/or larger stack volumes are strategies that can minimize errors depending on the structures of interest.

## Discussion

Recent developments in large-scale 3D imaging have created a bottleneck in automated image analysis. We have created an easy-to-use AI tool that is specifically suitable for biologists without expertise in programming. It closes the gap between large scale image acquisition and 3D visualization and allows the quantification of anatomical traits. Anatomics MLT provides the ability to extract quantitative data from files that may be tens of GB or larger with high precision. Statistical data on numbers, volumes, and volume fractions of specific organelles, cytoplasm to vacuole ratios, cell wall surface areas that are in contact with intercellular spaces versus neighboring cells, etc., can now be gathered comparatively easily. We foresee that Anatomics MLT will greatly facilitate the extraction of statistical data from different phenotypes at a precision that was previously impossible without enormous efforts. It will also provide the basis for inclusion of anatomical data into cell and tissue atlases and unltimately into holistic organismal models (Knoblauch and Peters, 2023).

Of fundamental importance in achieving good quality data is the embedding of the tissue samples. We optimized several protocols for plant tissues using *Arabidopsis* roots and leaves, as *Arabidopsis thalianais* still is the most commonly used model plant. In general, the optimization strategies we followed also apply to other plants, but it is trivial that plant species differ in their anatomy, metabolites, osmotic characteristics, turgor, etc. Those differences have an impact on the fixation and embedding procedures. Wavy cell walls, deflated vesicles and vacuoles, etc. are indicators that protocols need adjustments. We recommend working with experienced electron microscopists, ideally experts who are familiar with working on plant tissues.

The quantification of anatomical traits is of fundamental importance for recent efforts in crop improvement and the generation of cell– and tissue atlases (Lee *et al*., 2023; Tolleter *et al*., 2024). Anatomics MLT will provide a useful tool for gathering large-scale input data. A first example of the utility of Anatomics MLT, in this case for the quantification of chloroplast volumes, has recently been published (Knoblauch *et al*., 2024).

We provide Anatomics MLT as a community tool free of charge. The code is freely available as open-source, and we will welcome further additions and improvements by community members.

## Material and Methods

Whole *Arabidopsis* plate plants (age 6 to 30 days) were carefully lifted from the agar and **i**mmersed in room temperature fixative as specified in the protocol (Table 1). Microwave fixation was carried out in a BioWave Pro (Ted Pella, Redding CA). Block preparation for SBFSEM followed Lippens *et al*. (2019) except that approximately 30 nm of gold was deposited on the block to increase conductivity.

Imaging was performed in a Thermofisher Volumescope (Hillsboro, OR) either in high vacuum mode for conductive root tips or low vacuum for vacuolated plant tissue such as leaves. Imaging parameters varied depending on imaging conditions. For single energy imaging, pixel sizes ranged from 6 nm to 20 nm in the xy and from 40 nm to 100 nm in the z direction (z = slice depth for single energy imaging). Beam energy was kept at 1.78 KeV for high vacuum and 2.00 KeV for low vacuum imaging, with a beam current between 100 pA and 400 pA. Dwell times were adjusted to accommodate specific imaging requirements and ranged from 1 µs to 3 µs. Backscatter detection was collected with a T1 detector for high vacuum and a VS-DBS detector for low vacuum.

3D stacks were aligned, match contrasted, and Gaussian filtered with Amira Avizo software (Thermo Fisher, Waltham, MA).

### Reagent Preparation

Shelf lifes reported below are based on own experience and are only recommendations.

### Stock Solutions

**Cacodylate buffer** – 0.5 M stock buffer is prepared by adding 10.7 g of Cacodylic Acid Sodium Salt trihydrate to 90 ml ddH2O. The pH of the solution is adjusted to 7.2 with 10 N HCl and water is added to a final volume of 100 ml. The buffer can be stored at 4°C for up to 3 months.

**Calcium Chloride** – make 50 ml of a 0.2 M solution by adding 1.11 g CaCl_2_ powder to 50 ml of ddH_2_O. Can be stored at 4°C for up to 2 months.

**Glutaraldehyde** – 50 % vol/vol ampoules are opened using an ampoule cracker. The contents can be transferred to a scintillation vial and stored at 4°C for future use.

**Aspartic Acid** – 0.03 M aspartic acid is made by adding 0.4 g L-aspartice acid to 85 mL ddH_2_O. Adjust the pH to 5.5 with 1 N potassium hydroxide. Bring volume to 100 ml with ddH_2_O. Can be stored for up to 1 month at room temp. Check pH before each use.

**Fixative** –To make 30 ml of fixative, add 2.4 ml 50% glutaraldehyde, 0.3 ml of 0.2 M CaCl_2_, 1.5 ml of 0.5M cacodylate buffer to 25.8 ml ddH_2_O. This can be stored at 4°C for 3 to 4 weeks.

**Post stain 1** – Combine 0.375 g potassium ferrocyanide, 1.25 mL 0.5 M cacodylate buffer, 0.5 mL 0.1 M CaCl_2_ and 10.5 mL ddH_2_O.

**Post stain 2** – Combine 0.375 g 6-K hexacyanide, 1.25 mL 0.5 M cacodylate buffer, 0.5 mL 0.1 M CaCl_2_ and 10.5 mL ddH_2_O.

**Ligand** – 0.5% Thiocarbohydrazide is made by adding 0.1 g TCH to 10 ml H_2_O. Put into a 70°C oven for 60 min; the solution is swirled every 10 min. Once dissolved, take out of oven and add 10 ml H_2_O. Allow cooling to room temp and filter through a 0.2 µm syringe filter just prior to use. This solution must be made fresh prior to use each time.

### Stains

**Uranyl acetate** – add 1 g uranyl acetate to 50 ml H_2_O in a brown mixing bottle. Stir until dissolved. Filter twice through a new 0.2 µm syringe filter each time. Can be stored at room temp for about 1 month in a brown glass bottle.

**Walton’s Lead Acetate** – add 0.132 g lead nitrate to 20 ml 0.03 M aspartic acid (pH 5.5) in a glass bottle. Swirl until dissolved. Put in 60°C oven for 30 minutes. Flakey or cloudy solutions need to be replaced. Can be stored for up to 2 weeks at room temp.

### Resin

**Hard Spurrs resin without accelerator** – add 82 g ERL, 19 g DER, 118 g NSA to a large 500 ml Nalgene bottle with a screw on cap. Shake the solution for about 2 min. Divide resin into 6 50 ml Nalgene bottles with screw tops. Let this rest for 2 h at room temp then place into a –20°C freezer. Can be stored for up to 3 months at –20°C.

**Hard Spurrs resin with accelerator**– add 82 g ERL, 19 g DER, 118 g NSA and 2 g DMAE to a large 500 ml Nalgene bottle with a screw on cap. Shake the solution for about 2 min. Divide resin into 6 50 ml Nalgene bottles with screw tops. Let this rest for 2 h at room temp then place into a –20°C freezer. Can be stored for up to 3 months at –20°C.

## Acknowledgements

We thank the Franceschi Microscopy and Imaging Center at WSU for technical support and Dr. Daniel Mullendore for technical support and the testing of embedding protocols. This work was funded by grants NSF-PGRP 1940827 and NSF IOS 2318280 (MK), NSF-IIS-1553528 (AHG), NSF-IIS-2239688 (DW).

## Author Contributions

The work was conceptualized by MK, DW, HP, WSP, HHK, and AHG. Coding was performed by AB, YJ, and ZL under guidance of DW. Protocol testing and development was done by JK, BW and MK. The manuscript was written by MK and WSP with input from all authors.

## Appendix Please note this appendix is provided online at git hub together with the software. It is provided here for reviewing purposes only and will not appear in print

### Hardware

To process large 3D datasets, a sufficiently powerful computer is required. Ideally a computer cluster can be utilized. The reconstructions presented in the main text were done on an Intel Xeon w7-2475X, 2.59 GHz machine with 256 GB RAM and an NVIDIA RTX A6000 GPU. Training using 71 labels of statoliths at 500,000 iterations with a window size of 5, 501, 501 took about 24 h. Because of the smaller window size of 5, 129, 129 required for mitochondria (which are smaller than statoliths), the processing time with otherwise identical parameters was only 10 h.

Python supports NVIDIA graphics cards (GPUs). CUDA is a support software for NVIDIA GPUs. In order for Anatomics to make use of the GPU, a compatible CUDA version needs to be installed, which is dependent on the type of GPU that is used. Therefore, a CUDA version installed by default does not always work. When starting Anatomics MLT by typing “python gui.py” in the Anaconda prompt, the appearing lines should state:

*CUDA is available: True*

*Using Cuda Device* [name of your GPU])

If “*Cuda is available: False*” appears, a different CUDA version needs to be installed.

To check what CUDA version and driver are installed, type “NVIDIA-smi” into the command prompt. For more information on GPU requirements for python, visit https://pytorch.org/get-started/locally/

The following documents to install and test Anatomics MLT can be found at: https://github.com/PytorchConnectomics/Anatomics-MLT.

### Installation

– Since the Program requires “The PyTorch Connectomics package” which was mainly developed on Linux machines with NVIDIA GPUs, we recommend using **Linux** or **Windows** to ensure the compatibility of the latest features with your system. The instructions below are for **WINDOWS**.
– Install Miniconda https://docs.conda.io/en/latest/miniconda.html) following the instructions provided at the webpage.
– Open the “Anaconda Prompt”. You should be able to find this in the Windows start menu with your other programs. Either search for it, or look in the folder most likely called “Anaconda 3 (64-bit)”. Another way to find it is by clicking the start menu / press the Windows key, start typing miniconda, and select “Anaconda Prompt (Miniconda3)”
– Install the program using the commands below. Copy one line at a time by highlighting the line with the cursor. Press CTRL+C, or right click and select copy. Then run the command by pasting it into the Anaconda terminal (either using CTRL+V or right clicking and click paste). After you hit paste, the installation process should start. This may take a while. After installation, the line “Completely finished with installation” will appear. Copy the next line and continue the installation. Some systems allow copying and pasting all lines at once and the installation will run automatically. However, this does not always work.

### Commands to be executed for installation

cd Documents

conda create ––name plantTorch python=3.8.11 –y

conda activate plantTorch

conda install git –y

git clone https://github.com/PytorchConnectomics/Anatomics-MLT.git

cd Anatomics-MLT

conda install pytorch==2.0.1 torchvision==0.15.2 torchaudio==2.0.2 pytorch-cuda=11.8 –c pytorch –c nvidia

conda install cudatoolkit=11.8 –c pytorch

conda install h5py

git clone https://github.com/ajbrookhouse/pytorch_connectomics

cd pytorch_connectomics

pip install ––editable.

cd..

pip install open3d

pip install scikit-image

pip install paramiko

pip install pygubu

pip install pandas

pip install plyer

pip install ttkthemes

pip install connected-components-3d

conda install –c conda-forge imagecodecs –y

pip install neuroglancer

echo Completely finished with installation. Please run the program by typing ‘python gui.py’

– If the program does not open automatically, type “python gui.py” in the Anaconda terminal (usually this step should be completed automatically by the previous command copy section).

The main program should now be visible on your screen:

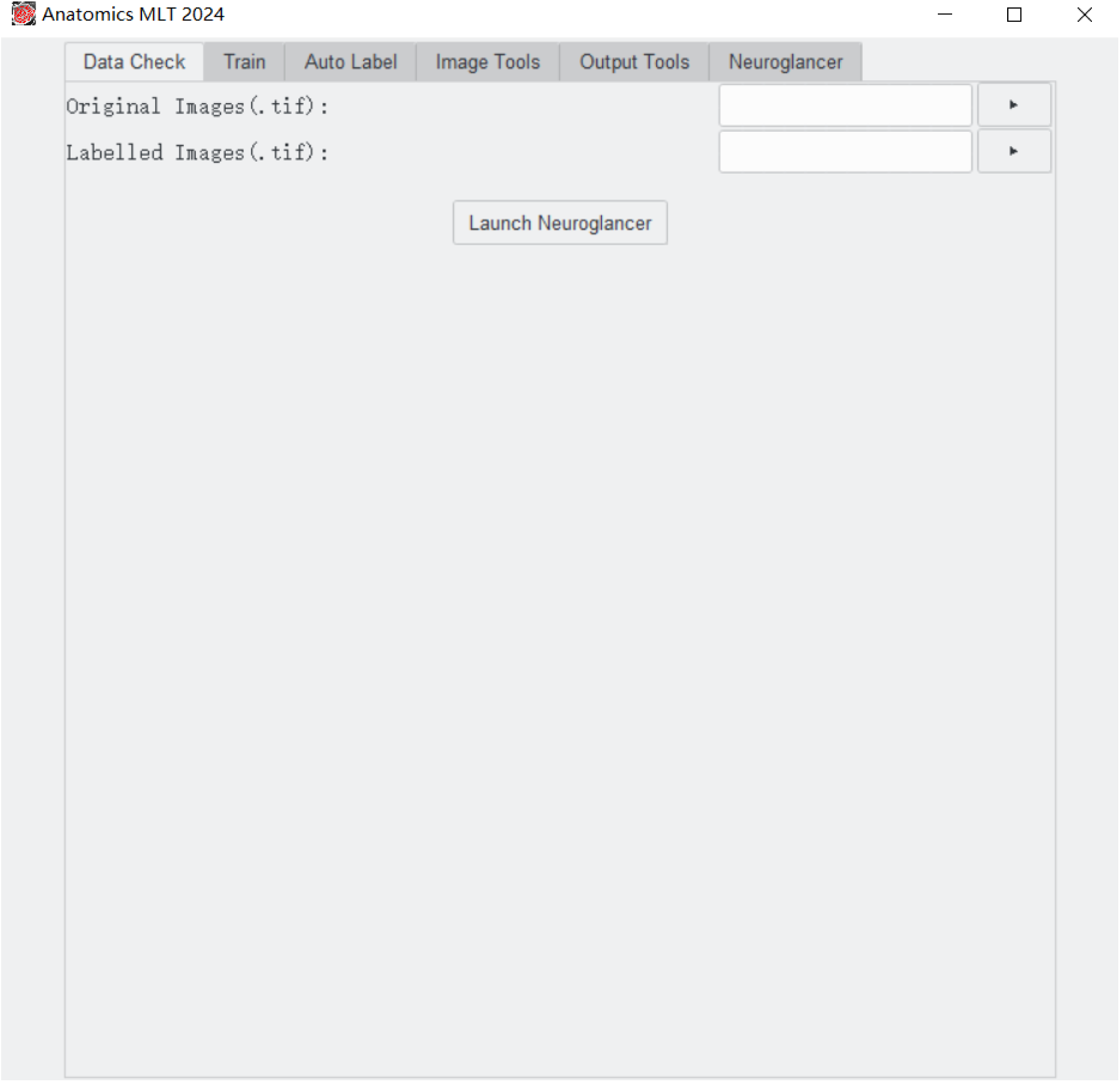

Updating Anatomics

Open miniconda by clicking start, typing miniconda, and selecting “Anaconda Prompt (Miniconda3)”. Then type the following:

cd Documents

cd Anatomics-MLT

git pull

If an error is shown when trying to update, please type ‘git reset ––hard’. Then use the command “git pull” to update the program.

### Uninstalling

If you need to uninstall the program for some reason (one reason could be getting a fresh install), close miniconda and delete the complete Anatomics-MLT folder. Then open miniconda and type the following:

conda deactivate plantTorch [only needed if your miniconda prompt lines start with (plantTorch).

conda env remove –n plantTorch –y

Now, all libraries used for the project will be uninstalled.

If you no longer need miniconda for other programs, feel free to uninstall it like any other windows program.

### QuickStart Guide

After installation of Anatomics MLT 2024, a test training may be accomplished by following the instructions below.

This tutorial presents an example of a semantic 3D segmentation.

Open Program

Open miniconda by clicking start, typing miniconda, and selecting “Anaconda Prompt (Miniconda3)”. Then type the following:

cd Documents

cd Anatomics-MLT

conda activate plantTorch

python gui.py

The main program should now be visible on your screen within a few seconds (if “python gui.py” does not work, try “python3 gui.py”). The window should look like this:

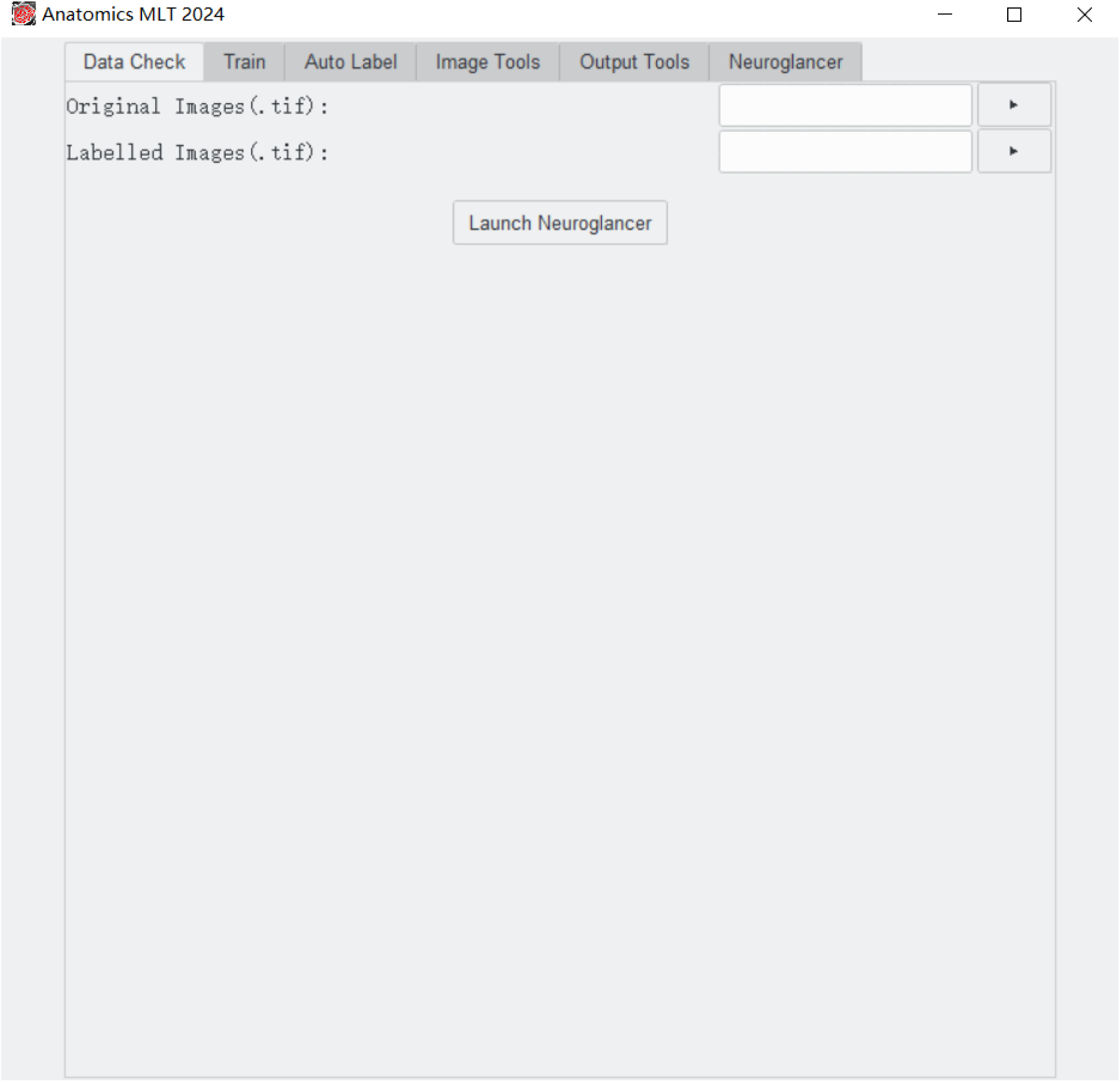

### Training Images and Labels

The first step is to obtain a training set of images, and a training label set. Below is an example of a training image:

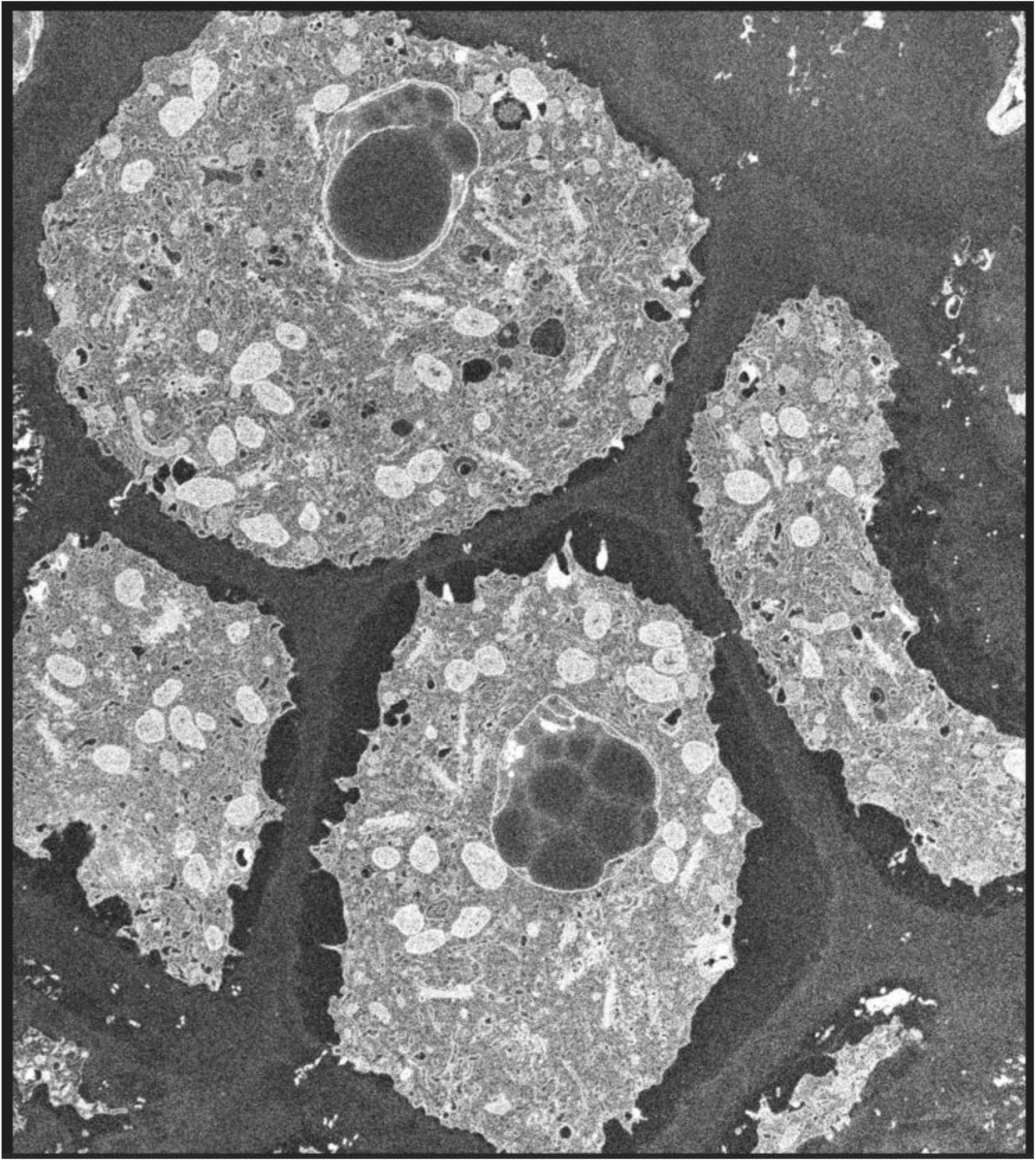

Below is an example of a training label:

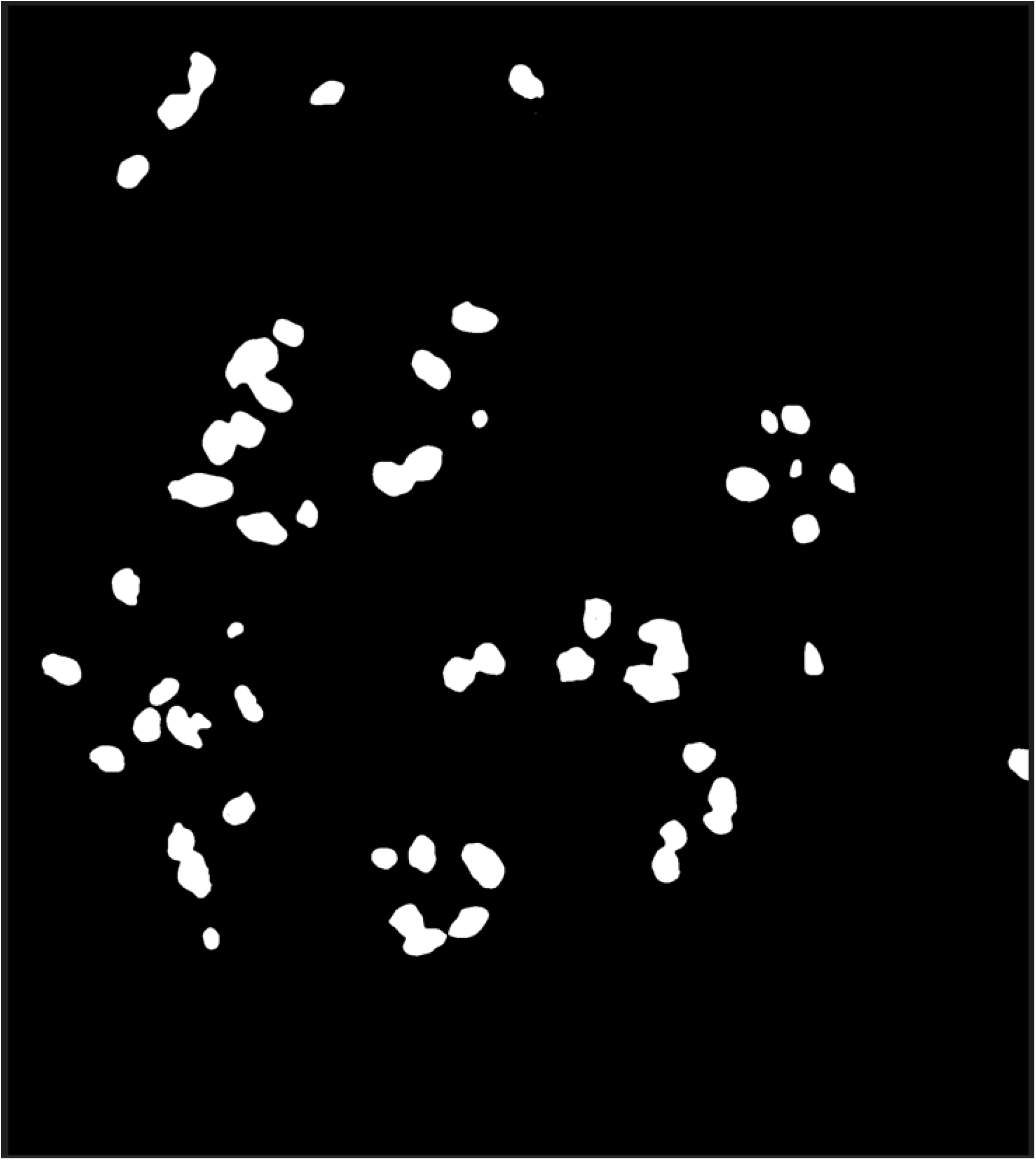

When using the models or algorithms in this program, it is important that image stacks are combined into a single stacked .tif file.

Because a semantic segmentation will be processed, the background is black and all instances are white.

### Data Check

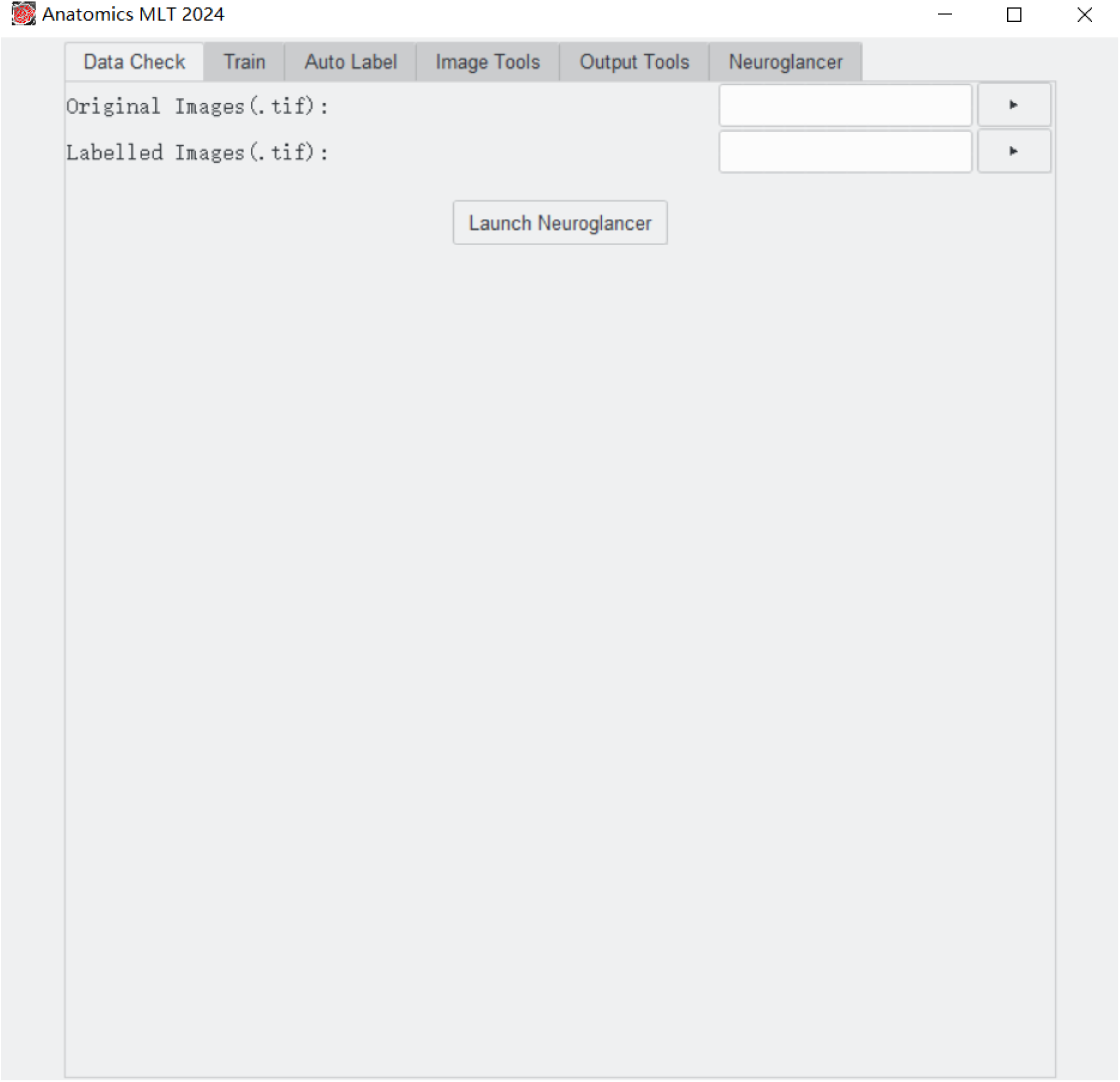

After producing a training dataset, the data format can be checked for accuracy in Anatomics MLT in the “Data Check” tab. Select the original image data stack in the upper box and the label stack that was generated from the image stack in the lower box. Click “Launch Neuroglancer”. A blue link will appear after a few seconds. Click the link to view the data in your browser. After the Neuroglancer is open, there will be four sub-windows. Each represents a different perspective (e.g. xy axis view; xz axis view; etc). We recommend using the view window to the lower right. In the top right corner of this sub window is a small button to enlarge the window. You may scroll through the z-layers of the image stack.

Put the cursor on the image. In the top left corner (next to “labels”) a number will display the grayscale value of the pixel defined by the cursor position. This number must be “0” if the cursor is positioned over the background. If the cursor is moved into an area where a specific organelle is labeled, this number must be larger than “0”. A number larger than “0” means the pixel is part of a label.

To avoid unnecessarily large file sizes, the preferred data format for label images and stacks is 8-bit grayscale or color. Only if more than 255 individual occurrences of the structure(s) of interest are present, a 16-bit (or higher) format should be used.

For semantic labels, the background should have the color value “0” and will be shown as black. All targets should have the same color (typically white). For example, if the image has 5 different mitochondria, the background should be black, and each mitochondrion will be labeled in white.

In case of instance labeling, the background color value must be “0”, and values for individual labeled areas (e.g. individual organelles) will have N different values larger than “0” (N stands for the number of different instances). For example, if the image has 5 different mitochondria, the background should be black, and each mitochondrion will be labeled in a different color (e.g. blue, white, red, green, orange) to distinguish between each instance.

### Training

To show the training window, click the “Train” tab on the toolbar on top.

In the following, we will discuss a training example based on datasets provided in the “ExampleData/” folder in the github repository.

The window should look like this:

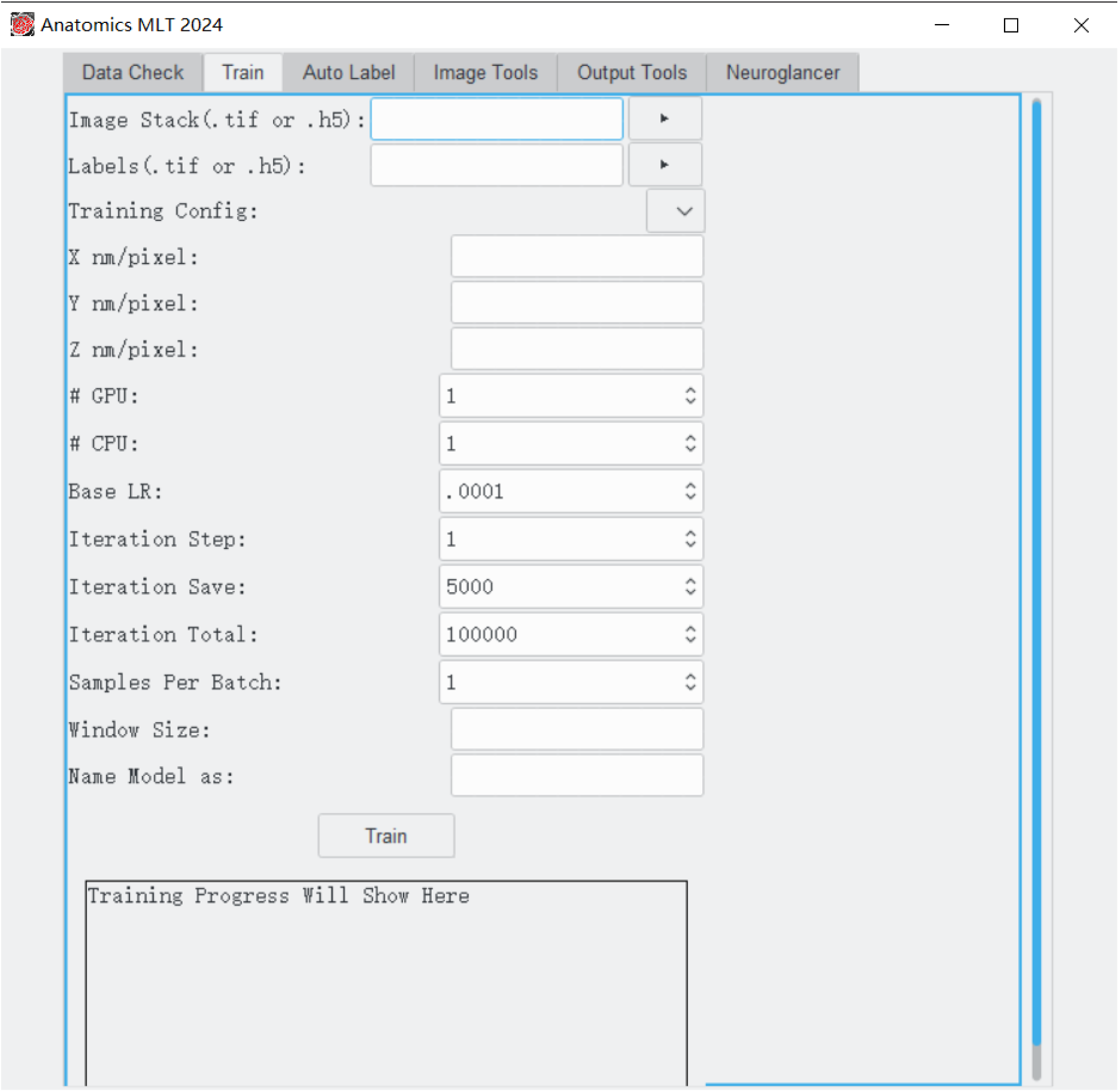

– Next to “Image Stack (.tif or .h5)”, click the triangle button to the right, and select “ExampleData/images 25-43_8-bit.tif”
– Next to “Labels (.tif or .h5)”, click the triangle button to the right and select “ExampleData/labels 25-43_8-bit adjusted 2.tif”
– Select the Training Config to be “Semantic3D.yaml”
– The example stack was created at x 10 = nm, y = 10 nm and z = 40 nm. Please type the appropriate numbers in the boxes labeled X, Y, and Z nm/pixel.
– In the text box labeled “Iteration Total”, enter 10000

Note: If you want to modify these boxes that have a default number input, please avoid using commas. For example, use 10000 not 10,000.

– In the text box labeled “Window Size:”, enter 3,129,129
– In the text box labeled “Name:”, give the model a name, for example: “Tutorial Network”
– Click the train button near the bottom. Information should start appearing in the text box. This process may take a long time depending on the capabilities of the computer. The text box will provide information on the Iteration number and an expected time to training completion. The training is complete when the last line reads “Rank: None. Device: cuda. Process is finished!”

### Automatic Labelling

Now that the model is trained, automatic labelling can be attempted. Click the tab “Auto-Label” near the top. The window should now look like this:

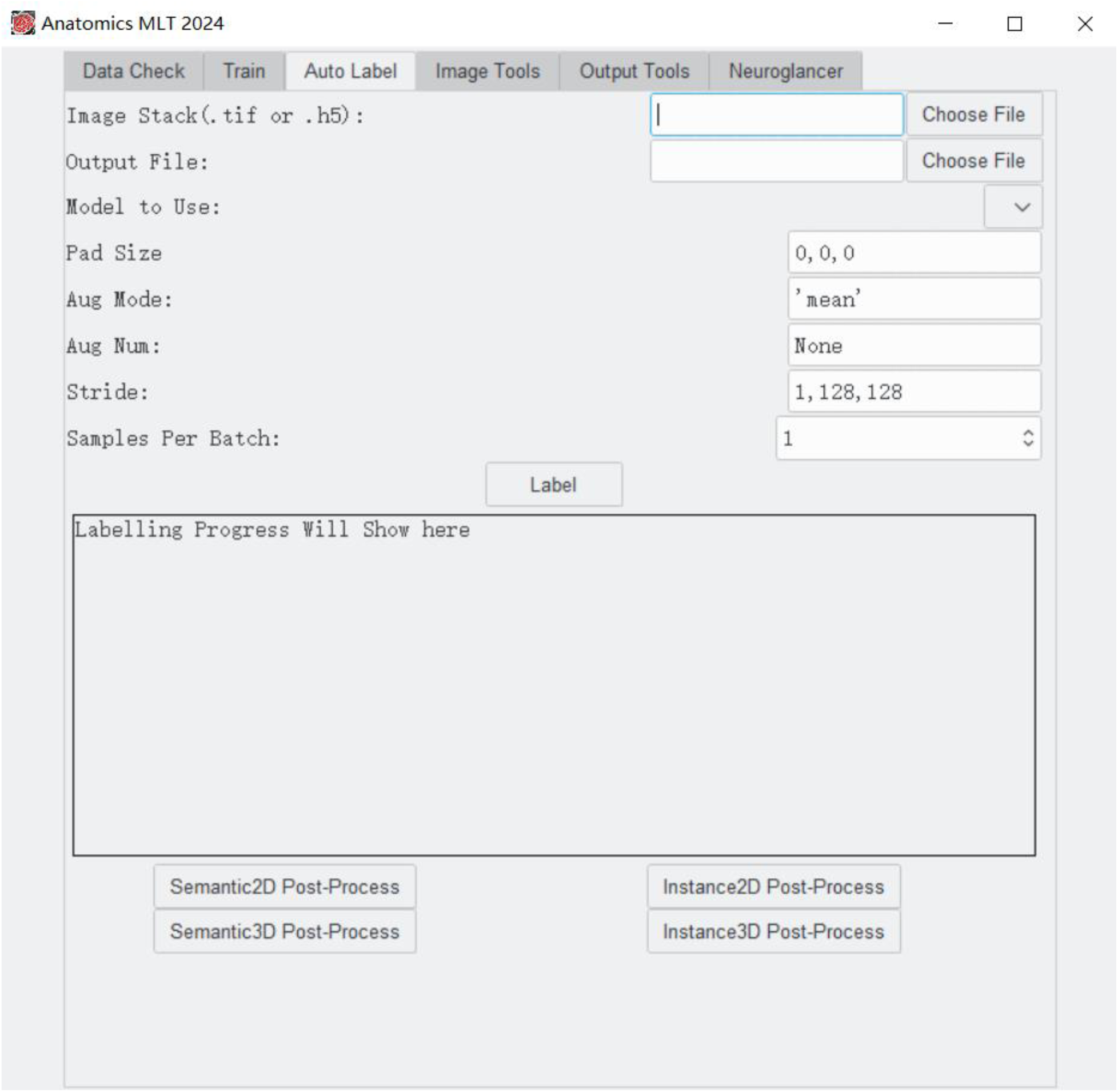

– In the “Image Stack (.tif)” selector, click the “Choose File” button, and in the dialog that pops up, select ExampleData/images 25-43_8-bit.tif

Note, it is not good practice to test the accuracy of a model on the same data that it was trained on. However, we will use the same training images here for illustration purposes.

– Click the “Choose File” button for “Output File:” and save the file as “ExampleData/myFirstPrediction”.
– Click the selection box to the right of “Model to Use:” and select “Tutorial Network”.
– Click label. This can also take a while but should be significantly shorter than the training. The output text box will print lines with information on “progress: {number}/{number} batches, total time {the estimated time to completion}. When the prediction is finished, the last line will read “Rank: None. Device: cuda. Process is finished!”
– Click Semantic post-process. This will post-process the initial output file and generate the final data output file, which will have the extension h5_s_out (instance post processing will generate a file with the extension h5_i_out).

Note: If your input and output files are not in the same folder, an error may show in the Anaconda prompt. You will need to reselect the file name under “chose file” and click “yes” when the question “file already exists. Do you want to overwrite” appears.

### Get Sample Stats

Now that the prediction is done, you can use the Anatomics to obtain statistics for the sample. Click the tab “Output Tools” near the top of the program. The window should now look like this:

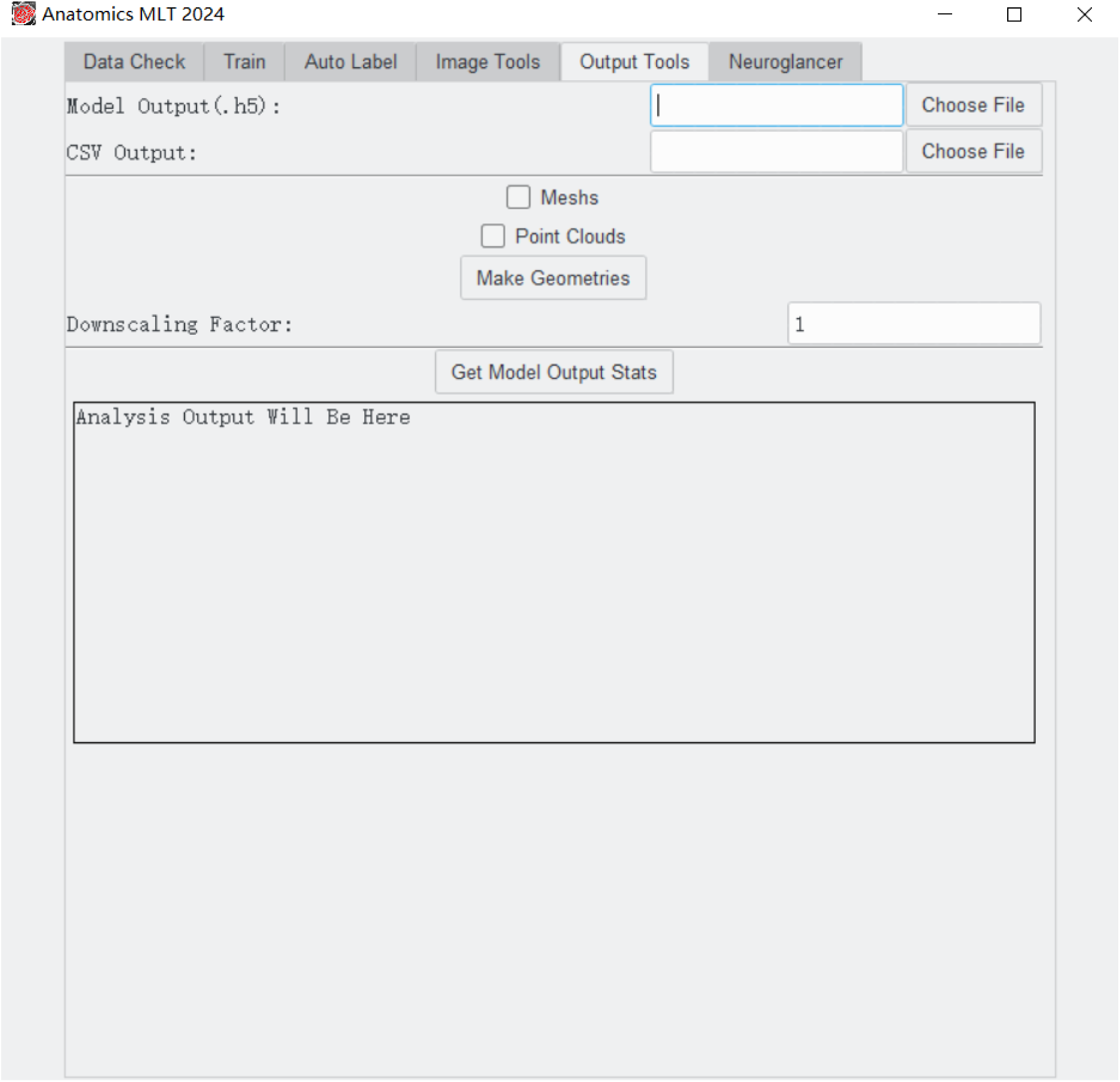

– In the file chooser labelled “Model Output (.h5)”, click on “Choose File”, and select “ExampleData/myFirstPrediction.h5” (if post processing was used the file will have the name “ ExampleData/myFirstPrediction.h5_s_out”)
– In the file chooser labelled “CSV Output”, click on “Choose File” and select a file name for the CSV file with output statistics.
– Click on the button named “Get Model Output Stats”. The program will show the min, max, mean, median, standard deviation, sum, and count of auto-labelled instances (mitochondria) in the sample. It will also generate an Excel (.CSV) file in your designated path.

Please note, with only 10,000 iterations for the training, the data will not be very accurate but sufficient to learn about the general process.

### Visualize

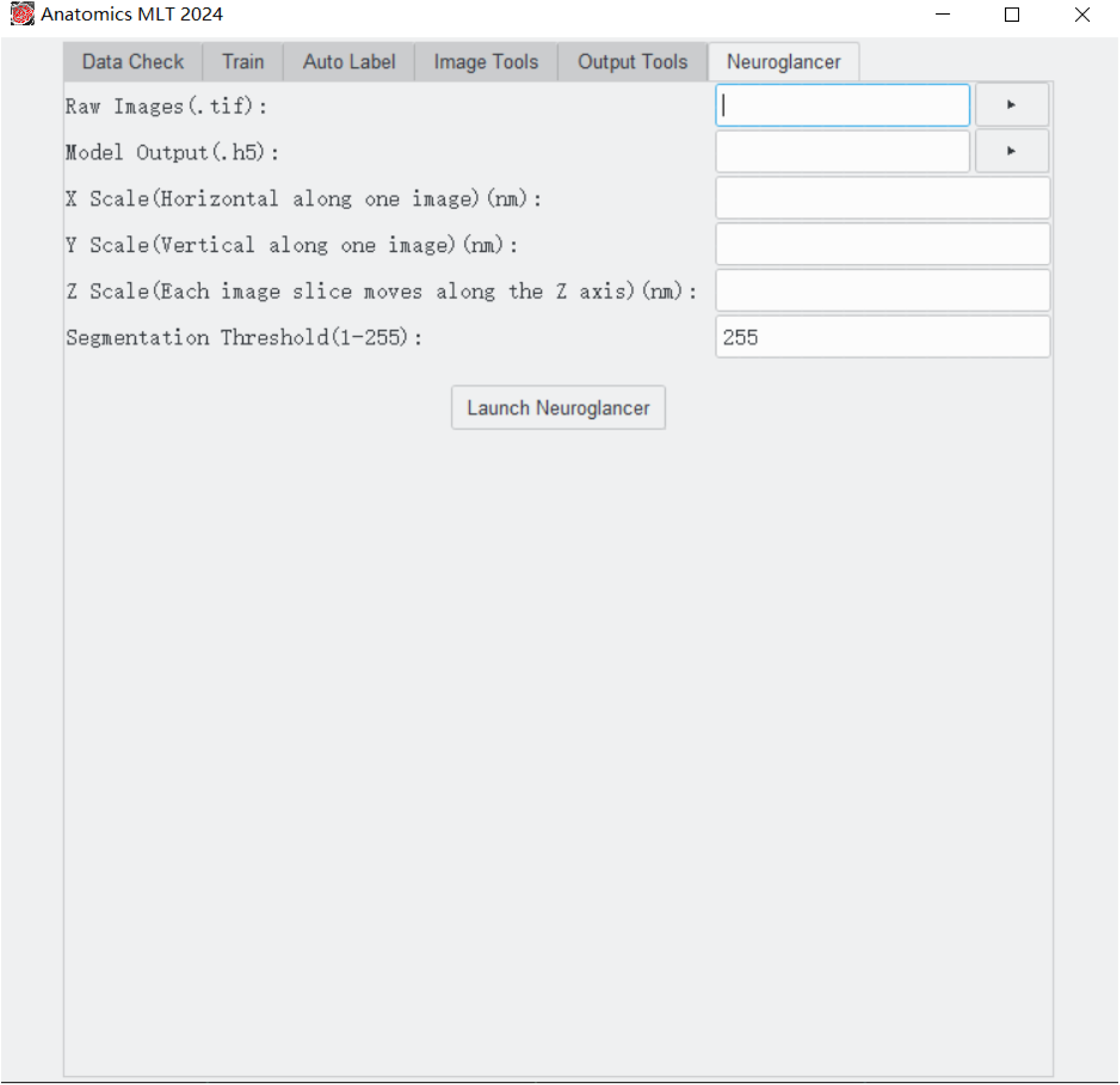

– Click the Neuroglancer tab.
– Select the Raw Image and Model Output by typing the name or using the interactive button.
– Enter the scales; in our case, z: 40; x: 10; y: 10.
– Segmentation Threshold; an 8-bit image has 256 grayscales or colors (0-255). Thresholding is a cutoff value that eliminates everything above this value. For example, if a value of 100 is chosen, only pixel with the value 0-100 will be shown. Keeping the value at 255, all grayscale or color values will appear in the final image.
– Click the “Launch Neuroglancer” to launch the visualization work. Once the visualization is ready, a blue link will show up in the software window. You can either click the link or copy and paste it to our browser to view the result.

### Datasets

– To create a 3D stack from individual images, all images should be in one folder and the images need to be named in consecutive order (e.g.0001.tif, 0002.tif, 0003.tif, etc.). The number represents the individual image’s location along the z axis. It is important that each file name has at least one leading 0, to ensure that the program orders them properly. A prefix is fine before the numbers in the filename, but it must be the same for all images. All images in the folder / stack must have the same dimensions and spatial resolution.

### The ImageTools screen

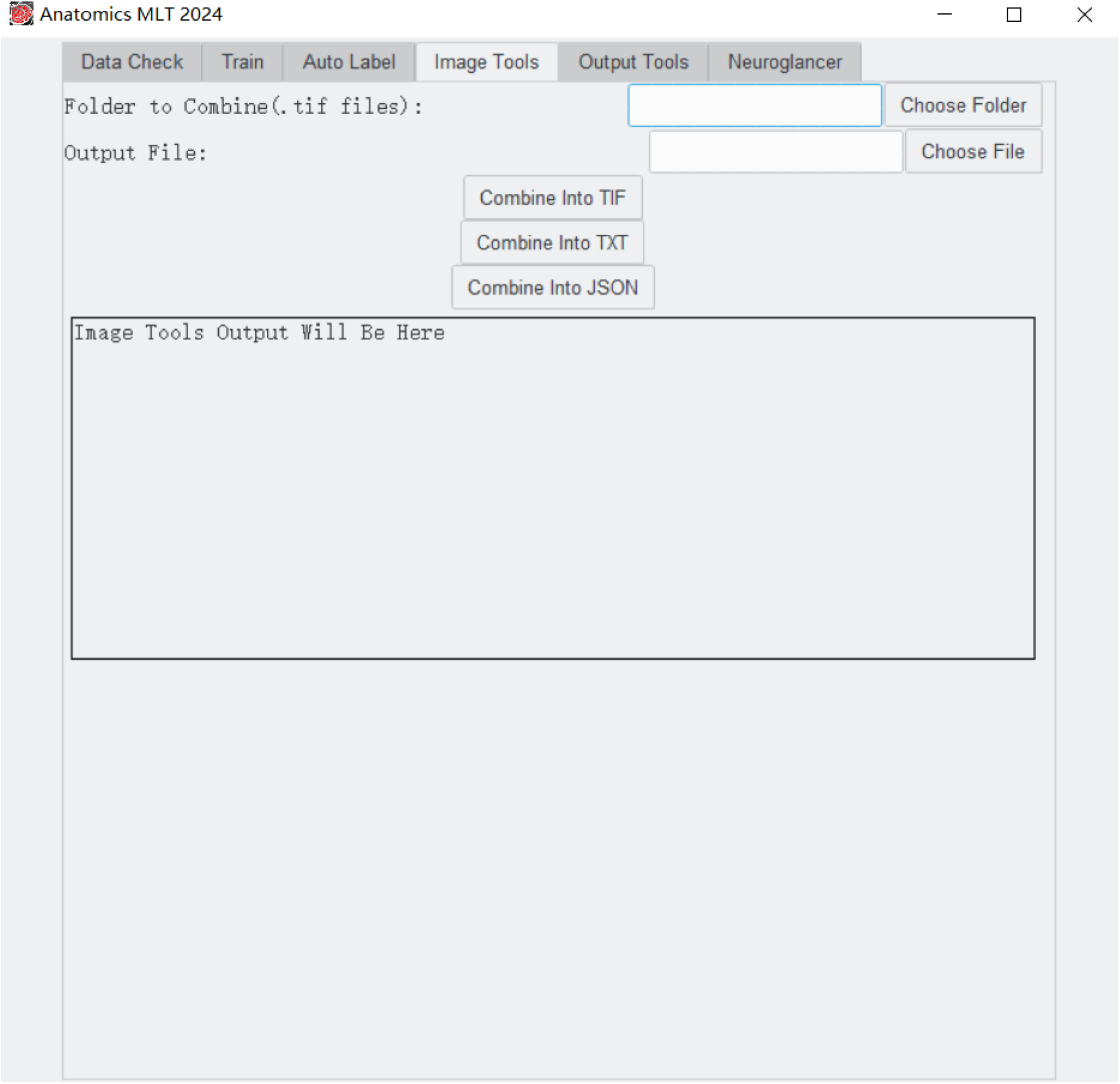

To create a single file for either Training or AutoLabelling, you must fill out the following fields:

– Folder To Combine: Click the “Choose Folder” button to pick the folder containing all images to be combined.
– Output Filename: Click the “Choose File” button and pick the location and name of your output file.

Next, click one (or more) of the “Combine Into” buttons.

– Combine into TIF: combines the dataset into a single .tif image stack. The 3D tiff file can be opened in most image software packages. However, tiff files are large compared to other formats.
– Combine into TXT: 2D only. This creates a .txt file for each individual image.
– Combine into JSON: .json files can process an arbitrarily large 3D dataset, and do not take up much space. They also allow the software to load smaller parts of the dataset at a time. Json files are suitable for processing extremely large datasets.

Note, both the TXT and JSON files contain the locations of the original images. This information is lost when moving the files.

### Detailed Description of Parameters

#### General Information

Semantic vs. Instance Segmentation

##### Semantic Segmentation

Semantic segmentation teaches the machine learning model to classify each pixel of the sample as one of several classes.

Pros:

– Can handle detecting different types of organelles at one time with the same model.
– Faster / requires less processing

Cons:

– The model does not differentiate between different instances of the same organelle. For example, if all mitochondria are labelled as “1”, the model does not understand the difference between the different mitochondria in the sample, it only understands whether a pixel belongs to mitochondria or not. Well separated mitochondria will still appear as individual instances, but mitochondria that touch each other will appear as one.

##### Instance Segmentation

Instance segmentation teaches the model to learn what pixels belong to one class, and their boundaries. In this way, it can differentiate two different organelles of the same type, even if they are touching each other.

Pros:

– The network itself learns how to differentiate between different instances of the same organelle.

Cons:

– Can only analyze one type of organelle at one time. For example, you can train the model to label chloroplasts or plasmodesmata, but not both at the same time. If instance segmentation for different structures is required, training two separate models is necessary.

##### Filetypes

There are several different filetypes that are used to store data in this program such as

.tif, .h5, .yaml, .json, and .csv files.

1. ”.tif” or “.tiff” files are used to store multi page images. These can be used to represent the 3D images that this program works with. It achieves 3D by stacking multiple 2D images together. However, please be aware that a large .tif will require a large computer memory space (RAM).
2. ”.h5” files, are like “.tif” files, but more versatile. “.h5” files can store arrays of any arbitrary dimension / size. They’re efficient to store large arrays or data and processing.
3. ”.yaml” files are used to store model configuration data/hyperparameters to define different models used by the program. When entering parameters in the interface, parameters will be passed to complete the yaml files. Once it is complete, the complete yaml file will be used by the algorithm to train the model.
4. ”.json” files can be used to make stacked/tiled datasets. However, it is not efficient for storing image data.
5. ”.csv” files are used to store statistical data. A CSV file will be generated once you click the “Get Model Output Stats” button. Then you may use Excel or other spreadsheet software to open, view and process.

#### Training

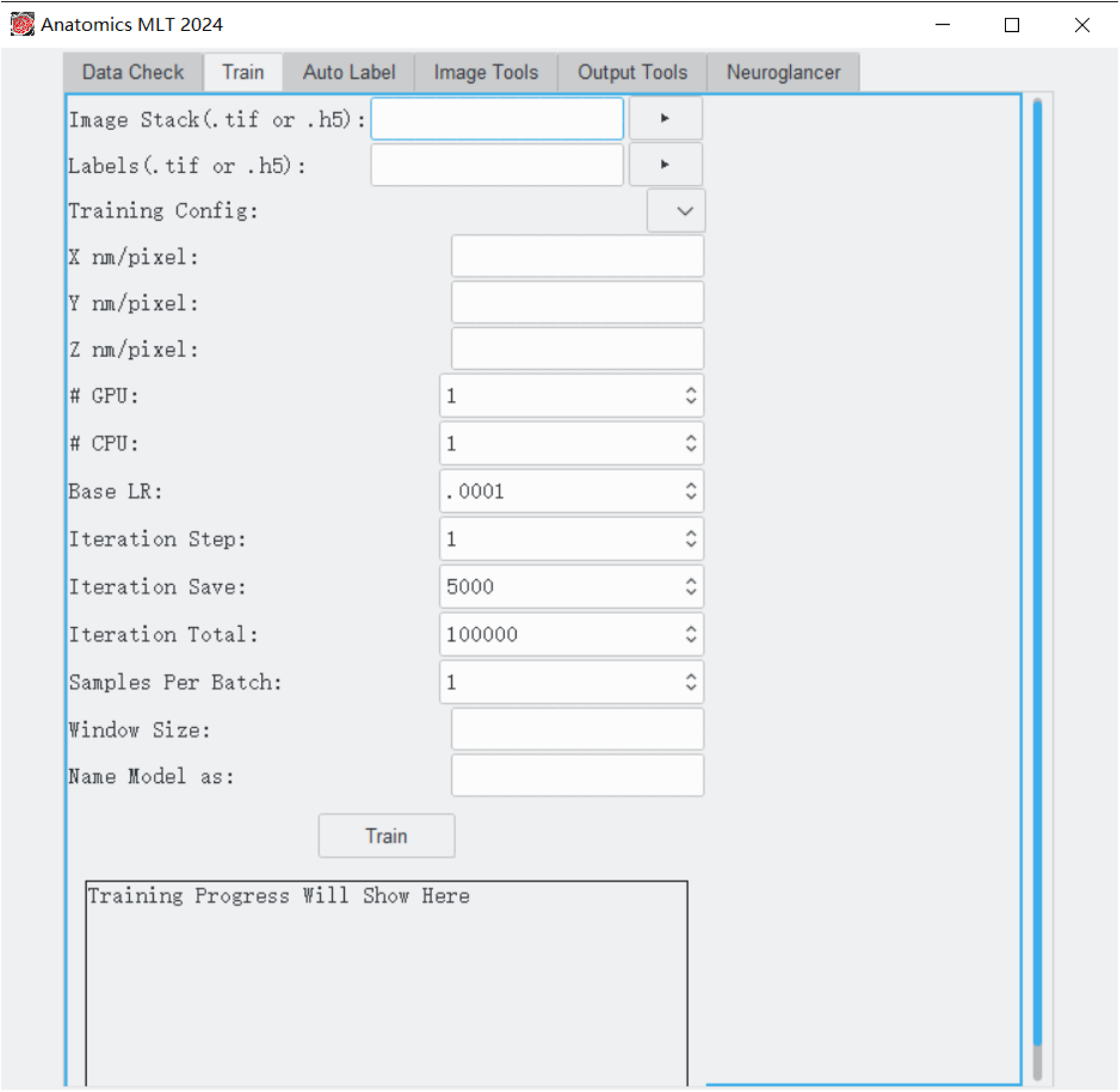

– Image Stack: Image stacks are the raw images that were originally generated on the microscope. The images should be very similar to the types of images that you want to process and may be images from the same 3D volume that will be used for the auto-label process (e.g. use images 1-50 to generate labels and the training stack and use images 50-1000 for the analysis).
– Labels: Labels are essentially a mask that tells the Anatomics MLT where the structures of interest are. More details about what labels represent can be found in the FAQ section.
– Training Config: Choose which type of model you want to train. By default, this program comes with Instance3D, Instance2D, Semantic2D, and Semantic3D.
– X nm/pixel: Provide spatial resolution for the image. In many cases the x, y, z resolutions are different (e.g. 10 nm X, 10 nm Y, 40 nm Z). If labels are not generated from the same stack that will be used for auto labelling, the training files should have the same x, y, z pixel dimensions.
– Y nm/pixel: Same as X, but for the y direction (vertical along one image slice)
– Z nm/pixel: Same as X, but for the z direction (through the stack of images, perpendicular to one image slice)
– GPU: The number of Graphic Processing Units (GPUs) to be used for the training. Most likely the answer will be 1, however if you have more (e.g. in a computer cluster) the number can be increased. While the program can run on the CPU only, an appropriate GPU is highly recommended to process large data sets.
– CPU: The number of Central Processing Units (CPUs) to be used for this training. The default is 1. Your system probably has 4, 6, or 8 in total to use if you want to increase this number. Do not increase this number above the number of CPU cores your computer has as this will cause a slowdown. This number can be increased and should speed up some calculations, however CPU does not affect the speed of training a model nearly as much as the GPU does.
– Base LR: Learning Rate (LR) is a parameter that should be chosen every time a model is trained. Essentially, this changes how quickly the model changes its internal parameters each time it processes a subset of data. A higher learning rate causes the model to change faster. This can lead to the model learning faster / needing less training, however a higher learning rate can also lead to unstable training or cause the model to not fully optimize. We recommend a starting value of 0.0001
– Iteration Step: Number of iterations the program runs simultaneously (typically, 1).
– Iteration Save: The program will incrementally save your model as it trains. It does this every multiple of this number. For example, if 100000 iterations are used, and iteration save is 10000, the model would save at 10000, 20000, 30000, etc. It is recommended to save a few times during the training process, but too often will use unnecessary amounts of disk space. About 10-20 saves during the entire training process is sufficient. The number must be equal to or a fraction of “Iteration Total”.
– Iteration Total: The total number of training iterations. A higher number means the model trains over more data points. This is an important parameter to improve training quality, but it also linearly increases computation time. If too few steps are chosen, the model will not be accurate enough to properly identify structures of interest. If too many iterations are chosen the model may “overfit” the training data, meaning the model becomes too biased towards the training data and cannot perform well on test data. Typical iteration numbers for good training are 300000 to 500000.
– Samples Per Batch: How many data iterations to process at the same time. 1 means the software investigates one data point, and using this output it adjusts the parameters and moves on to the next data point. If 2 is chosen, two data points are processed simultaneously and then the network parameters are updated. This may lead to higher quality training but doubling “samples per batch” doubles computation time and may cause problems with the computer memory. Typical numbers are 1 to 4 but may be higher if a computer cluster is available.
– Window Size: The model or detector operates/trains in a small window or kernel. This window slides over the input data for training. It must have three numbers, separated by commas; these numbers define the size of the window or kernel. The numbers must be separated by commas (z, x, y). Typically, the window size is determined by the target that has been labeled. At a minimum, a label should cover an entire structure of interest. For example, if mitochondria are labeled a small window size such as 3,129,129 is sufficient because the mitochondria in our example stack are comparatively small. In the case of statoliths in our example, a window size of 3, 501, 501 is required because the statolith are much larger and have a diameter of approximately 300-500 pixel. It is not necessary to increase the window size beyond the size of the structure of interest as this will consume more RAM and lead to longer processing time.
– Name Model as: Provide a unique name that can be recognized for future auto-labelling processes.

### Auto-Labelling

To use the Auto-Labelling Feature, you must first train a model that can be chosen for the Auto-Label process.

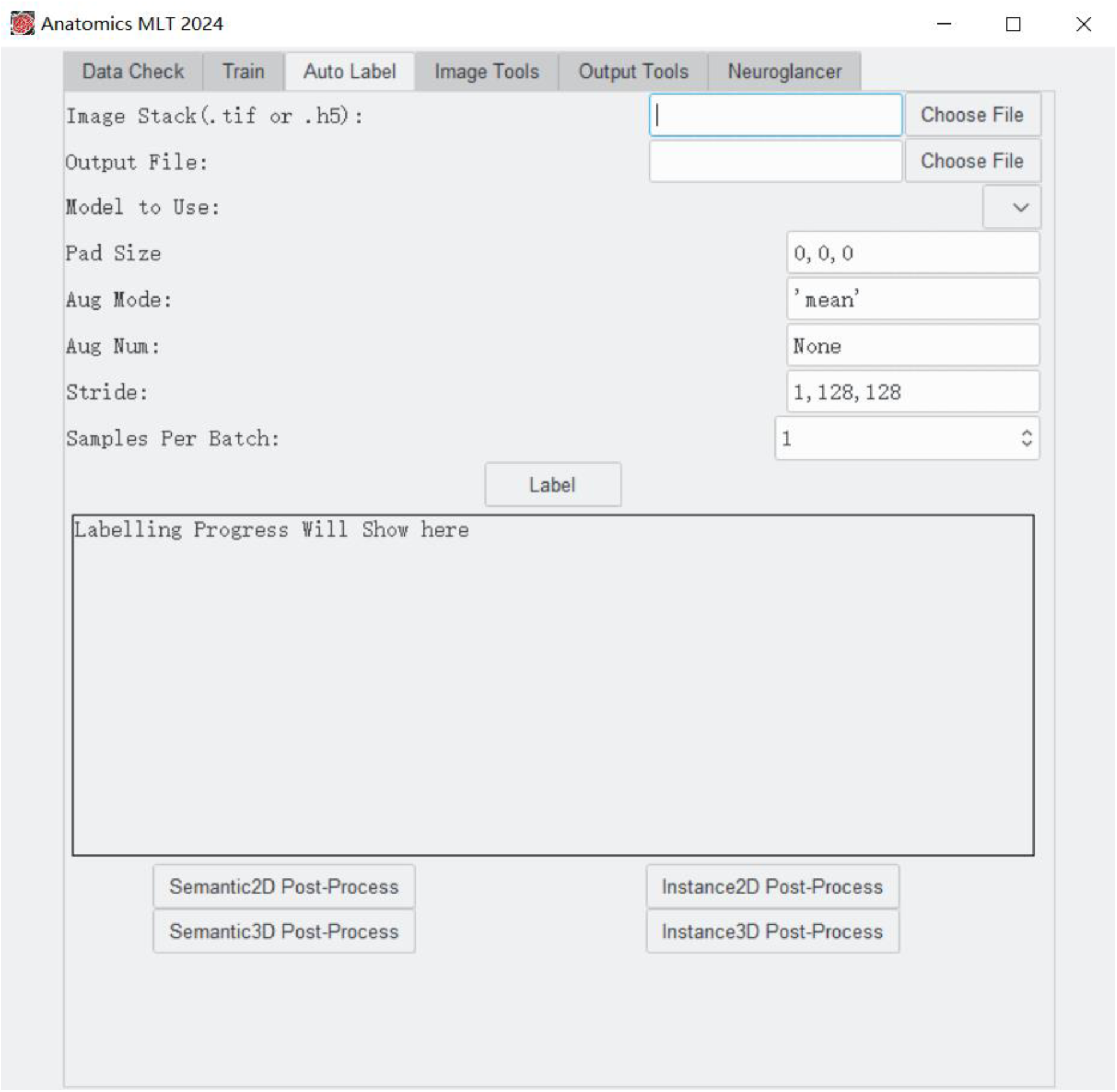

#### Detailed Description of Each Parameter

– Image Stack: This file is a 3D image stack of original images (usually) taken with a microscope that will be used for analysis.
– Output File: Provide an output file name (this is a new file to be created) that will contain the identified structures of interest and that will be used for statistics and visualization.
– Model To Use: Select the name of the model to be used. This is a file that was generated during the training using the labels highlighting the structures of interest. All training files will automatically appear in the drop-down menu when clicking the arrow.
– Pad Size: The model can add padding around the outside of a section that the model is currently processing. More info can be found at https://deepai.org/machine-learning-glossary-and-terms/padding.
– Aug Num: Augmentation number. Each input data volume can be transformed in multiple ways, e.g., flipping the xy dimension, and the model can be run on the modified data to obtain multiple results. We can specify the augmentation number to “4” or “8” to apply pre-defined transformations. “None” means that no transformation is applied.
– Aug Mode: Augmentation Mode. Given the multiple results from above, we can apply either “max” or “mean” to combine them into the final result. We would recommend using the ‘mean’ mode which empirically leads to better results in general.
– Stride: The model does not process the entire dataset at once. It works on small sections at a time. After it has finished processing a section, it moves on and processes the next section. The amount it moves after each iteration is the stride. We recommend using a stride that corresponds to the window size that was used during the training. Smaller strides than the window size will work, but bigger strides will result in non-processed sections.
– Samples Per Batch: How many data iterations to process at the same time. It has no implication for model quality (Auto-Labelling), other than that processing may be faster when the number is slightly increased. Higher numbers require better hardware.

Label: Clicking this button will start the labeling process.

The text box will indicate the labeling progress. Once finished it will indicate which post-processing step is needed. If a post processing step is required, please click the appropriate box below the text box (e.g. Semantic3D Post-Process…)

### Image Tools

Folder to Combine: Click “chose folder” and select the folder that contains all files you want to have combined into a single 3D stack. The folder may not contain other files

Output file: provide the path and name of the file that will be generated. Combine into Tif: Click this button to create a 3D tiff file.

Combine into TXT: Click this button to create a TXT file for each individual image. Combine into JSON: Click this button to create a 3D JSON file.

### Output Tools

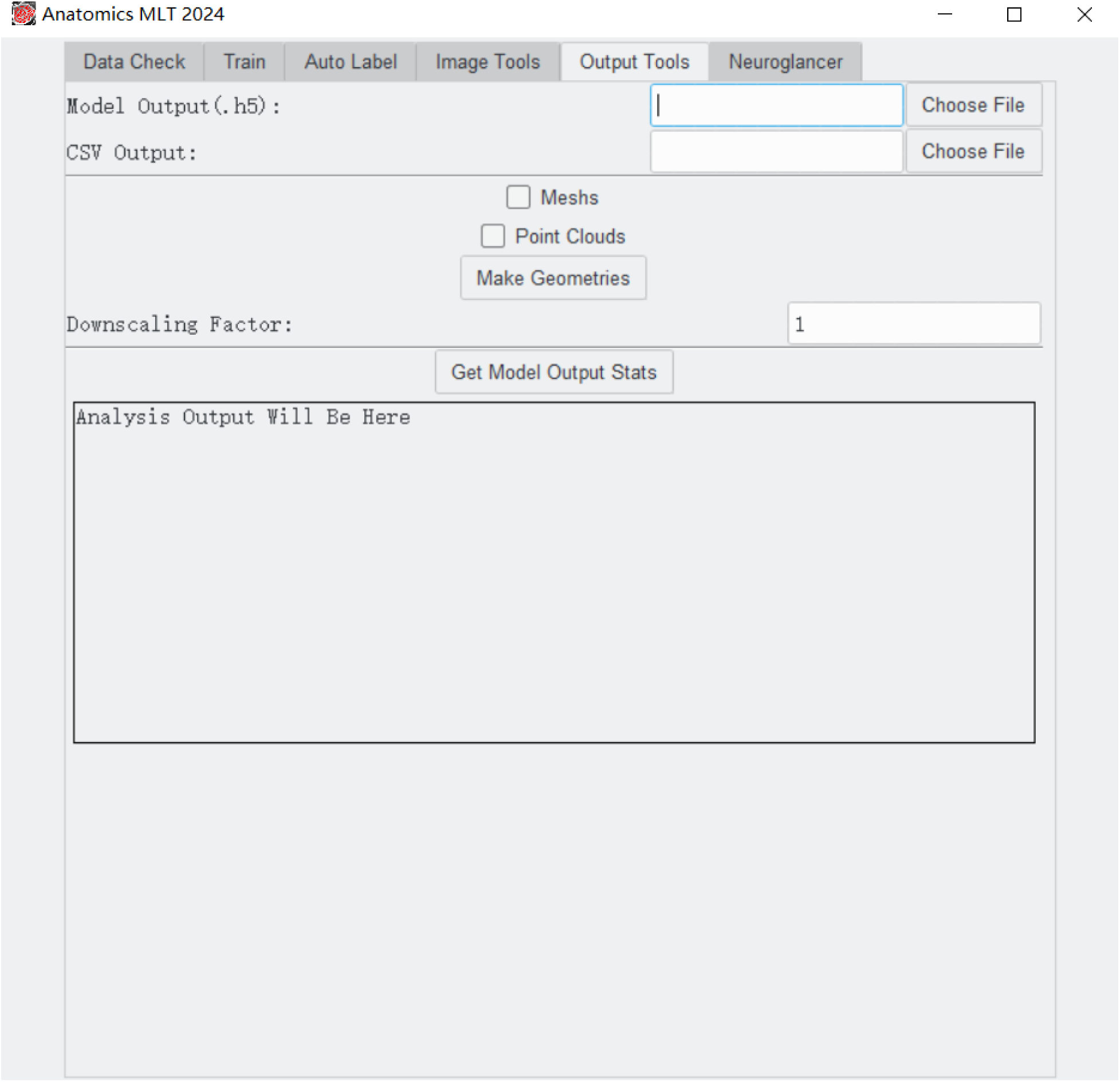

Model Output: Chose an .h5 (or h5_s_out, or h5_i_out if post-processing was done) file of interest that was generated during the auto-label process.

CSV Output: Provide a name for the statistical data output file to be generated.

Meshes: highlight this box if you want to receive a 3D mesh that can be imported into Blender, Amira, and other 3D image processing software.

Point Clouds: highlight this box if you want to receive a 3D point cloud that can be imported into Blender, Amira, and other 3D image processing software

Make Geometries: Click this button if you want to create a mesh or a point cloud or both. The program will generate .ply files for the meshes and point clouds that start with the name of the original .h5 file followed by .ply

If you want to create a downscaled sampled version of the geometry, type a downscaling factor into the Downscaling Factor field. This factor works on all three axes, for example if you use a downscaling factor of 2, all three axes will be halved in size and the total volume will be reduced by a factor of 8.

Get Model Output Stats: Click this button if you want to receive a CSV file containing statistical data. The stats will also be printed into the text box once they are calculated (this could take a while for semantic data but should be quick for instance data).

### Neuroglancer

This tab allows the use of Neuroglancer, a visualization package, to visualize the generated data.

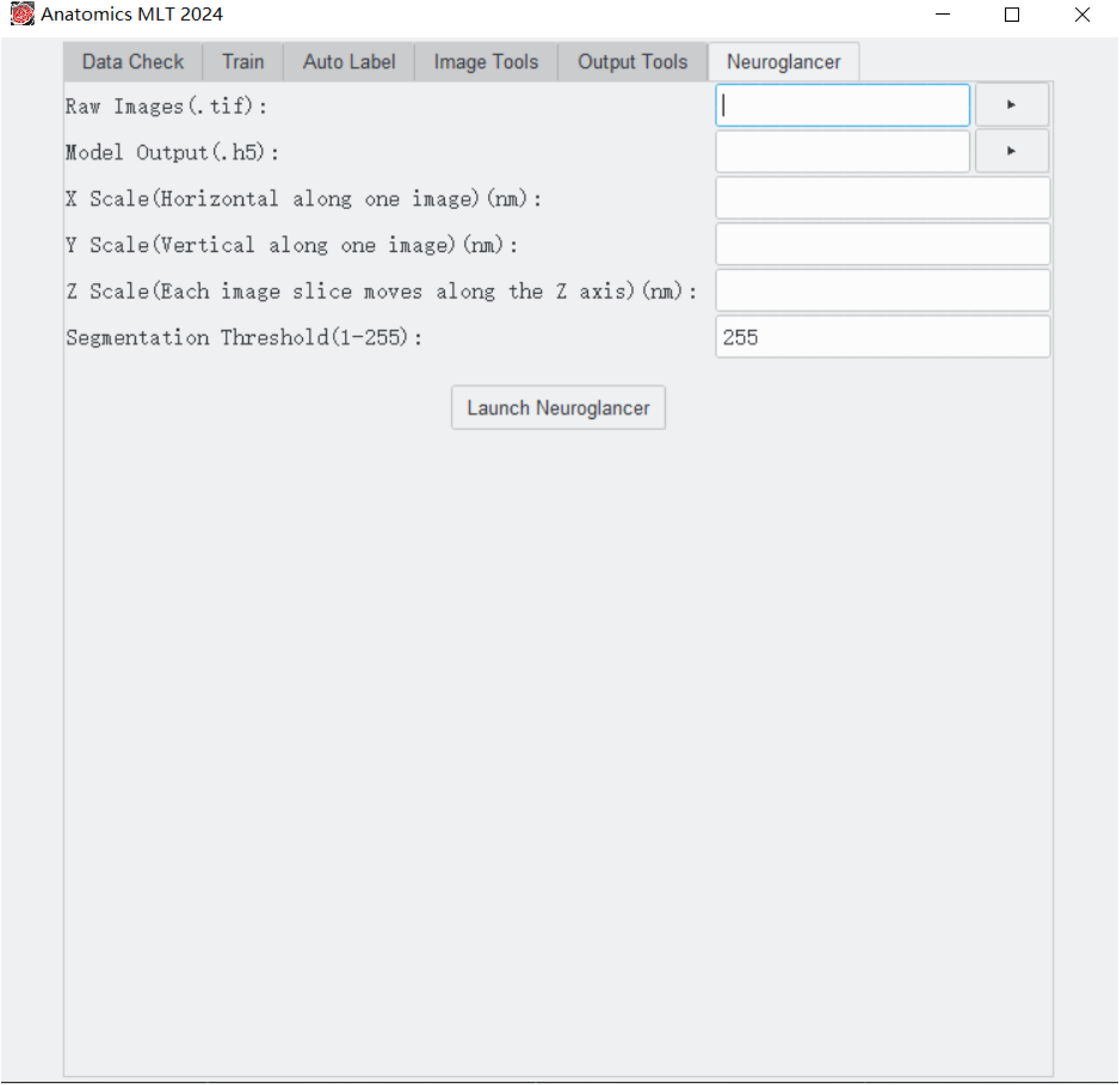

Raw Images: Select the image stack that was used for the auto-label process.

Model Output: Select the .h5 file that was generated during the auto-label process. If you used a semantic 3D model, the file would end with “.h5_s_out”; if you used an instance 3D model, the file would end with “.h5_i_out”.

X Scale, Y scale, Z scale: Enter the X scale, Y scale, and Z scale. Neuroglancer needs to know if there are non-square pixel dimensions.

Segmentation threshold: This is a number between 0 and 255 and can cut off certain grayscales. Use 255 to keep all data.

Click “Launch Neuroglancer”. Once the visualization is ready (which may take a while), a blue link will appear in the software window. Clicking the link will open the default browser and will display the reconstruction.

